# Physicochemical Targeting of Lipid Nanoparticles to the Lungs Induces Clotting: Mechanisms and Solutions

**DOI:** 10.1101/2023.07.21.550080

**Authors:** Serena Omo-Lamai, Marco E. Zamora, Manthan N. Patel, Jichuan Wu, Jia Nong, Zhicheng Wang, Alina Peshkova, Liam S. Chase, Eno-Obong Essien, Vladimir Muzykantov, Oscar Marcos-Contreras, Jacob W. Myerson, Jacob S. Brenner

## Abstract

Lipid nanoparticles (LNPs) have become the dominant drug delivery technology in industry, holding the promise to deliver RNA to up- or down-regulate any protein of interest. LNPs have been targeted to specific cell types or organs by physicochemical targeting, in which LNP’s lipid compositions are adjusted to find mixtures with the desired tropism. In a popular approach, physicochemical targeting is accomplished by formulating with charged lipids. Negatively charged lipids localize LNPs to the spleen, and positively charged lipids to the lungs. Here we found that lung-tropic LNPs employing cationic lipids induce massive thrombosis. We demonstrate that thrombosis is induced in the lungs and other organs, and greatly exacerbated by pre-existing inflammation. This clotting is induced by a variety of formulations with cationic lipids, including LNPs and non-LNP nanoparticles. The mechanism depends on the LNPs binding to fibrinogen and inducing platelet and thrombin activation. Based on these mechanisms, we engineered multiple solutions which enable positively charged LNPs to target the lungs while not inducing thrombosis. Our findings implicate thrombosis as a major barrier that blood erects against LNPs with cationic components and illustrate how physicochemical targeting approaches must be investigated early for risks and re-engineered with a careful understanding of biological mechanisms.

## Introduction

Since the first FDA approval of a solid lipid nanoparticle (LNP) in 2018, LNPs have rapidly become the preferred drug delivery platform of the biopharma industry, with the trend accelerating markedly after the COVID-19 LNP vaccines^1^. Targeting LNPs to specific organs or cell types is desirable for many applications and is generally accomplished by one of two methods. The older method is to conjugate LNPs to affinity moieties (e.g., antibodies) that bind to a known epitope on a target cell^2^. This method presents major challenges for scale-up manufacturability and immunogenicity, partially contributing to it never having produced an FDA-approved targeted LNP or liposome. The second, and newer, method for LNP targeting is to screen large numbers of LNP formulations for those that have fortuitous “physicochemical tropism” to a target organ. In physicochemical tropism/targeting, some (usually unknown) physical or chemical features of the LNP cause it to gain entry into particular cells. Over the last decade, the physicochemical approach has become the dominant method of achieving targeting in both academia and industry, because of the ease of *in vivo* screening and the highly efficient manufacturing process. As physicochemical tropism has been able to target many different organs and cell types, it has the potential to meet LNPs’ promise of being able to modulate any protein in any cell type or organ.

Physicochemical targeting is usually achieved by screening many lipids, typically varying each of the 4 major lipid classes used in LNPs: ionizable lipids, PEGylated lipids, cholesterol, and helper lipids (phospholipids, sphingolipids, etc). The mechanisms of targeting are rarely elucidated. As a notable exception, ApoE was implicated as the plasma protein that binds to the first FDA-approved LNP, patisiran, shuttling the LNPs to hepatocytes^3, 4^. While mechanisms of delivery generally aren’t elucidated, there have been some clear trends in physicochemical properties that correlate with organ distribution. For example, intravenously injected (IV) LNPs formulated with negatively charged lipids often accumulate in the spleen^5–9^, and LNPs formulated with positively charged lipids usually accumulate in the lungs^5, 7, 9–12^, as found independently by many labs and companies. These observations were, in a 2020 *Nature Nanotechnology* paper, distilled into a simpler and more elegant way of targeting LNPs^5^. Instead of using particular ratios of hard-to-synthesize ionizable lipids or exotic helper lipids, this seminal study showed it was possible add to a base formulation of LNPs either a negatively charged lipid to target the spleen or a positively charged lipid to target the lungs. This approach leads to democratization and reproducibility, since any lab or company can obtain such common lipids, rather than having to synthesize the particular ionizable lipid variants used in a particular lab’s screening.

However, while testing such promising physicochemically-targeted LNPs as therapeutics, we found that the LNPs with lung tropism induce a major unreported side effect: thrombosis. Studies of nanoparticle toxicities have largely focused on 2 of the 3 major defense systems of plasma: complement proteins and immunoglobulins^13–15^. However, the 3rd defense system, clotting, has often been neglected in recent years, even though it has the most deadly and acute consequences if improperly activated. Blood has evolved to actively clot in response to different foreign surfaces and the coagulation cascade is specifically known to activate in response to charged surfaces. An IV dose of nanoparticles exposes the blood to massive amounts of foreign surface area, but nanoparticles can also indirectly activate clotting by interacting with cells that affect clotting processes, such as endothelial cells, neutrophils, and platelets. Noting that the dominant nanomedicine targeting method now employs manipulation of nanoparticle charge, it is important to revisit the thrombotic side effects of nanomedicines^16–21^, with a focus on the new targeting approaches.

Here we show that lung-tropic LNPs with positively charged lipids induce massive thrombosis. This finding was quite general, as it held for LNPs with widely varying ionizable lipids, different positively charged lipids, and not just LNPs, but also liposomes. The lung-tropic LNPs induced profound pulmonary embolism when administered intravenously, induced stroke-like effects when administered via the carotid artery, and modified clotting processes in *ex vivo* blood. We have investigated detailed mechanisms of LNP-induced clotting, showing lung-tropic LNPs bind to the core clotting protein, fibrinogen, and cause aggregation and activation of platelets. Elucidating these detailed mechanisms of clotting enabled us to develop three solutions that may permit positively charged LNPs to maintain their excellent lung targeting while not inducing dangerous clotting: Pre-treatment with anticoagulants (direct thrombin inhibitors, but *not* the seemingly obvious choice of heparin), conjugation of the LNPs to direct thrombin inhibitors, or reduction of the LNP size.

## Main

### Lung-tropic LNPs with cationic lipids induce gross side effects in vivo

We fabricated physicochemically-targeted, lung-tropic LNPs by the previously published method of adding in a cationic lipid, DOTAP (1,2-dioleoyl-3-trimethylammonium-propane (chloride salt)). We initially synthesized these LNPs with the ionizable lipid cKK-E12 (herein referred to as +DOTAP LNPs, unless otherwise stated). This base +DOTAP LNP formulation consisted of the lipids DOTAP, cKK-E12, 1,2-dioleoyl-*sn*-glycero-3-phosphoethanolamine (DOPE), cholesterol, 1,2-dimyristoyl-rac-glycero-3-methoxypolyethylene glycol-2000 (DMG-PEG 2000) (50/25/5/18.5/1.5, mol/mol) and mRNA (lipid/mRNA ratio = 40/1, wt/wt). Control, non-lung-tropic LNPs were synthesized without DOTAP (Fig. 1a). There was no significant difference in the sizes of these LNPs (99.22 ± 4.27nm for -DOTAP LNPs and 102.47 ± 0.48nm for +DOTAP LNPs) (Fig. 1b). However, +DOTAP LNPs had a significantly higher zeta potential than -DOTAP LNPs (14.34mV vs −4.05mV) due to the addition of the cationic lipid (Fig 1c).

**Figure 1:**
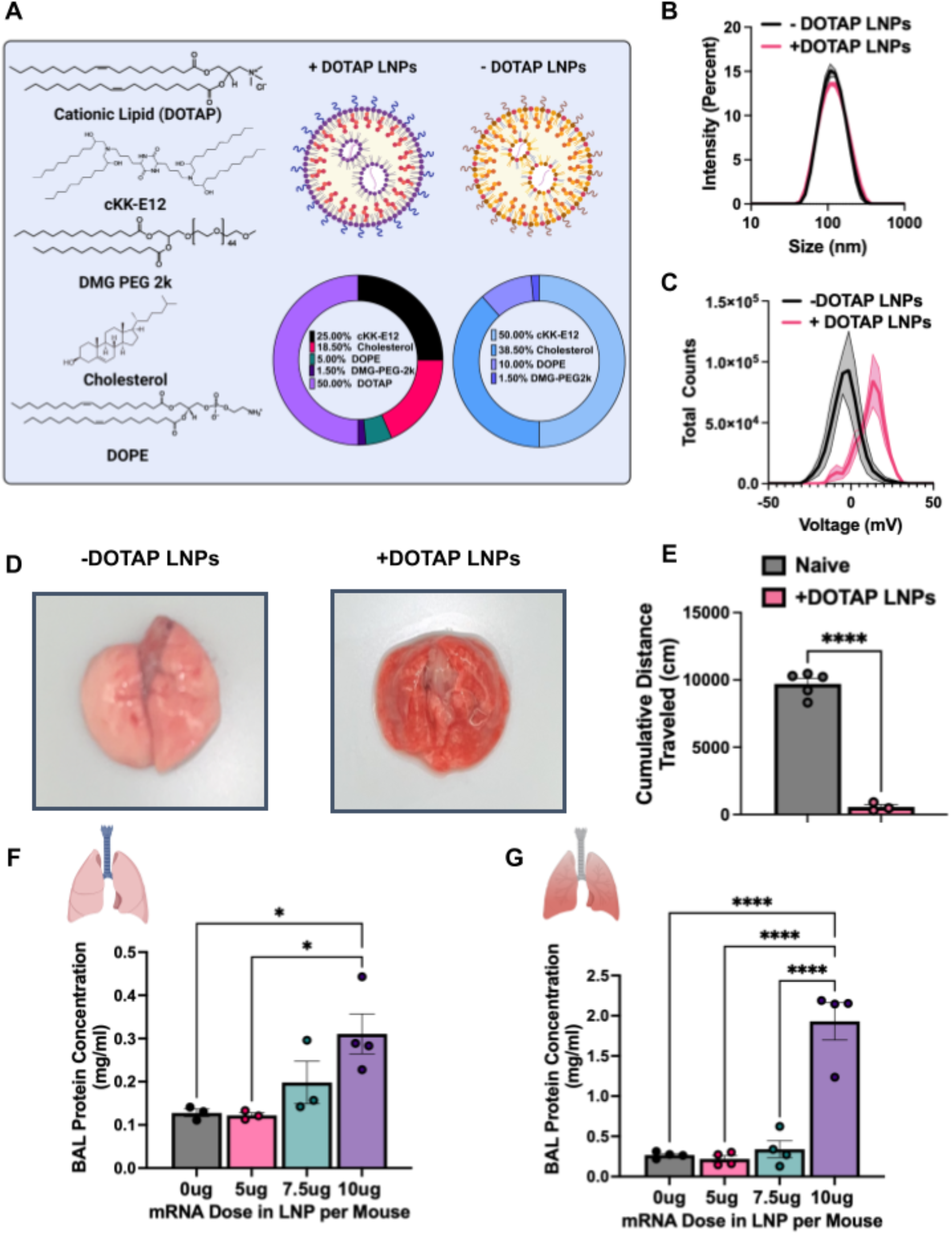
LNPs with physicochemical tropism to the lungs induce acute, severe side effects in mice. (A) Formulation of -DOTAP and +DOTAP LNPs. (B) Size distribution of - DOTAP and +DOTAP LNPs determined by dynamic light scattering (DLS) shows no significant difference in size of these LNPs (99.22 ± 4.27nm for -DOTAP LNPs and 102.47 ± 0.48nm for +DOTAP LNPs). (C) Zeta potential graphs of -DOTAP LNP versus +DOTAP LNP show that - DOTAP LNPs are slightly negatively charged at −4.05mV and +DOTAP LNPs have a surface charge of 14.34 mV. (D) Gross anatomical comparison of naïve mice intravenously (IV) injected with either -DOTAP LNPs or +DOTAP LNPs reveals severe lung hepatization (liver-like appearance) with +DOTAP LNP injected mice. LNPs were allowed to circulate for 30 minutes. (E) Comparison of cumulative distance traveled for 1 hour after injection of +DOTAP LNPs assessed by AI software, DeepLabCut, compared to naïve, uninjected mice reveals a significant reduction in total distance traveled, indicating lethargy in these mice. (F) Dose-response of the effect of +DOTAP LNP dose on protein concentration in bronchoalveolar lavage (BAL) fluid shows dose-dependent increase in BAL protein, a measure of endothelial barrier dysfunction. (G) This is further exacerbated in mice with pre-existing lung inflammation (nebulized LPS mice; note the 5-fold y-axis augmentation between F & G). Statistics: n=3-5 and data shown represents mean ± SEM; For (E), comparisons between groups were made using an unpaired t-test with Welch’s correction. For all other graphs, comparisons between groups were made using 1-way ANOVA with Tukey’s post-hoc test. *=p<0.05, ***=p<0.001, ****=p<0.0001.

We next intravenously (IV) injected the above LNPs into naive mice at a dose of 10µg of mRNA per mouse and sacrificed 30 minutes later for gross anatomical inspection. We observed that in mice that received +DOTAP LNPs, the lungs were strikingly red and firm, resembling “hepatization” (liver-like appearance) described by pathologists for human acute respiratory distress syndrome (ARDS) (Fig. 1d). We observed clear lethargy and sluggishness in +DOTAP LNP mice. Using the software DeepLabCut, we found that after a recording time interval of 1 hour post LNP injection, +DOTAP LNP mice walked a total distance that was >17-fold less than naïve mice (Fig 1e). We further assessed lung-specific toxicity by measuring the contents of the bronchoalveolar lavage (BAL) fluid 20 hours after IV +DOTAP LNP injection, which assesses lung capillary leak. In naive mice, there was a dose-dependent increase in both BAL protein and leukocyte count from 5µg to 10µg of mRNA (Fig. 1f and Supplementary Fig. 2).

Because lung-targeted LNPs will generally be given only to patients with lung pathology, we next investigated whether +DOTAP LNPs caused even more severe injury in mice with pre-existing lung inflammation. We utilized a mouse model of acute lung inflammation achieved by administering nebulized lipopolysaccharides (LPS). +DOTAP LNPs were IV injected 4 hours after LPS exposure, and mice were sacrificed 20 hours later. As in naïve mice, we saw a dose-dependent increase in BAL protein concentration from 5µg to 10µg of mRNA (Fig. 1g). However, in nebulized-LPS mice, there was an exacerbation of BAL edema at each dose tested. At the 10µg dose of mRNA, nebulized-LPS mice have 6-fold higher BAL edema compared to naïve mice injected with +DOTAP LNPs. This indicates that the toxicity of +DOTAP LNPs is amplified under pre-existing inflammatory conditions. Based on these data, a dose of 10µg of mRNA in LNPs was used for further in vivo studies unless otherwise stated. This is a therapeutically relevant dose for mRNA-LNP delivery and enabled us to adequately assess the mechanisms behind DOTAP LNP toxicity.

### Coagulation is triggered by LNPs and other nanoparticles with cationic lipids, across diverse formulations

Given the very acute nature of the side effects of +DOTAP LNPs, seen within a minute on the AI-measured distance-walked test, we hypothesized that the nanoparticles were activating a protein-based defense system in the blood. The three major plasma protein defense systems are the complement cascade, immunoglobulins, and the coagulation cascade. Since complement has been long implicated in nanoparticle defense^13–15^, and since immunoglobulins’ acute effects also lead to complement activation (via IgG-mediated activation of the classical pathway of complement), we first investigated if these LNPs activate the complement system. Surprisingly, we discovered that +DOTAP LNPs do not activate complement, as we detected no increase in the plasma concentration of C3a 30 minutes after +DOTAP LNP IV injection into naive mice (Supplementary Fig. 3, noting we have extensively shown C3a production induced by other nanoparticles^13, 14^)). Without C3a production, it is highly unlikely that complement activation, or acute immunoglobulin effects, mediate +DOTAP LNPs’ acute toxicity.

Therefore, we investigated whether the toxicity was produced by the coagulation cascade. To investigate this, we stained lung sections with Masson’s trichrome to look for evidence of clots. Lung histological samples show large clots both at the large vessel and capillary levels in naïve mice IV injected with +DOTAP LNPs, compared to mice injected with -DOTAP LNPs where there are no observable clots (Fig. 2a). This was a clear indication of thrombosis (clotting) due to +DOTAP LNPs. Notably, this effect is not restricted to the lungs since we also observed evidence of clotting in the brain when +DOTAP LNPs were injected via the carotid artery (Supplementary Fig. 4).

**Figure 2:**
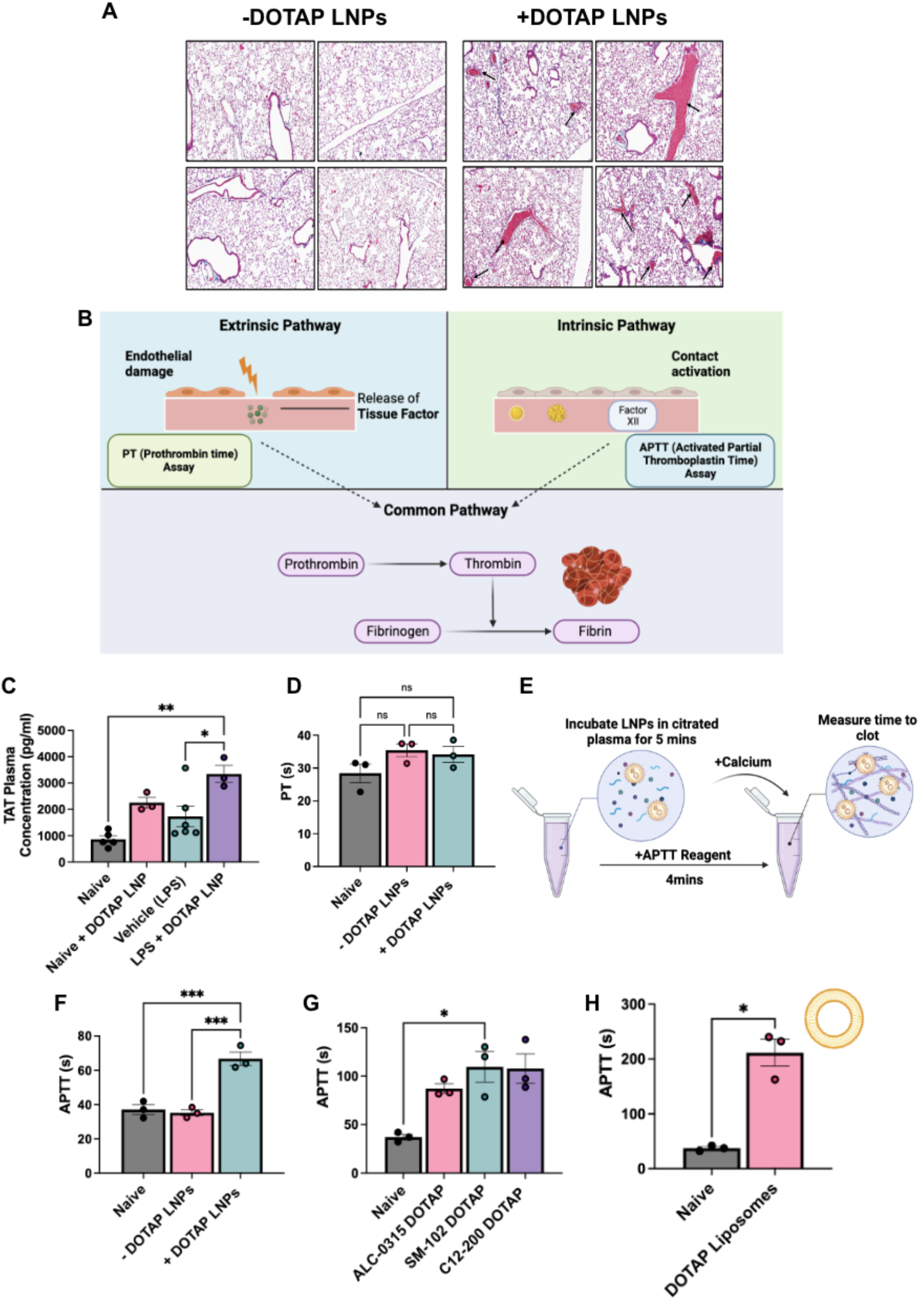
Coagulation is activated by positively-charged LNPs and other nanoparticles, across diverse formulations, both *in vivo* and in *ex vivo* whole blood. (A) +DOTAP LNPs induce large clots in the pulmonary arteries (black arrows). -DOTAP or +DOTAP LNPs were injected into healthy mice at a dose of 10ug of mRNA and 30 minutes after, lungs were harvested and prepared for histology (Masson’s Trichrome). -DOTAP LNPs do not show such clots. (B) Schematic of the two major coagulation pathways - intrinsic and extrinsic - which both converge into the common pathway. The extrinsic pathway is triggered by damage that occurs outside the blood vessel, exposing of tissue factor. The intrinsic pathway is initiated by factors within the blood vessel lumen. Both the intrinsic and extrinsic pathways converge to the common pathway, the conversion of prothrombin to thrombin - an enzyme that converts fibrinogen to fibrin. Fibrin forms the mesh-like framework of clots. The PT (Prothrombin Time) and APTT (Activated Partial Thromboplastin Time) are laboratory tests used to measure the time to clot through the extrinsic and intrinsic coagulation pathways, respectively. (C) *In vivo*, +DOTAP LNPs increase plasma levels of thrombin-antithrombin (TAT), a marker of recent clotting. The TAT increase is > 2-fold in naive mice, with an additional > 1.5-fold increase in mice with pre-existing lung inflammation (nebulized-LPS). +DOTAP LNPs were injected into naïve mice and 30 minutes post-injection, plasma was collected for TAT assay. (D) +DOTAP LNPs do not change PT time in vitro, showing that they do not induce coagulation through the extrinsic coagulation pathway. (E) Timeline and schematic of *in vitro* APTT measurements. (F) +DOTAP LNPs increase APTT, showing that they induce coagulation through the extrinsic pathway (G) +DOTAP LNPs increase APTT through the intrinsic pathway regardless of ionizable lipid identity, as ALC-0315, SM-102, and C12-200 DOTAP LNPs also show elevated APTT. (H) Positively charged, non-LNP nanoparticles, here liposomes containing DOTAP, also increase APTT time. Statistics: n=3-6 and data shown represents mean ± SEM; For (H), comparisons between groups were made using an unpaired t-test with Welch’s correction. For all other graphs, comparisons between groups were made using 1-way ANOVA with Tukey’s post-hoc test. *=p<0.05, ***=p<0.001, ****=p<0.0001.

We needed to determine which molecular pathways led to LNP-induced thrombosis. Coagulation can be divided into the intrinsic and extrinsic pathways (Fig. 2b). The extrinsic pathway is activated by trauma that damages the endothelium, leading to exposure of tissue factor which activates the coagulation protein Factor VII. The intrinsic pathway is initiated by factors within the blood vessel lumen, and has been shown to be triggered by charged surfaces (such as on foreign particles) or collagen, which trigger the activation of the coagulation protein Factor XII^22^. The extrinsic and intrinsic pathways converge to the common pathway which involves the conversion of prothrombin to thrombin - the enzyme that converts fibrinogen to fibrin. Fibrin forms the mesh-like framework of clots. The PT (Prothrombin Time) and APTT (Activated Partial Thromboplastin Time) are laboratory tests used to measure the time to clot through either the extrinsic or intrinsic pathways, respectively.

We measured thrombin-antithrombin (TAT) plasma levels in both untreated and nebulized-LPS injured mice. TAT is a stable complex formed after thrombin activation that serves as a marker of coagulation. We found that +DOTAP LNPs increase TAT levels by >2-fold in naive mice and by an additional >1.5-fold in nebulized-LPS injured mice (Fig. 2c). This confirmed that +DOTAP LNPs activate thrombin (the common pathway of coagulation), and this is exacerbated in pre-existing inflammatory conditions, probably as a result of thrombo-inflammation^23^. To determine if +DOTAP LNP-activated coagulation was employing the intrinsic or extrinsic pathway, we measured PT and APTT after doping LNPs into plasma in vitro. While +DOTAP LNPs did not change the PT (Fig. 2d), they led to a significant increase in APTT (Fig. 2e, f). This indicates that +DOTAP LNPs activate the intrinsic coagulation pathway, thus depleting intrinsic pathway proteins and prolonging the APTT.

We formulated liposomes and other LNPs with DOTAP to see if the coagulation side effects were a generalizable result of co-formulation with cationic lipids. We formulated LNPs with various ionizable lipids (C12-200, ALC-0315, SM-102; Supplementary Fig. 1 and Supplementary Table 1) and found that +DOTAP LNPs increase APTT, regardless of the ionizable lipid type (Fig. 2g). We used DOTMA, rather than DOTAP, as our cationic lipid and observed the same effect on APTT (Supplementary Fig. 5). Finally, adding DOTAP to liposomes, rather than LNPs, also results in increased APTT (Fig. 2h and Supplementary Fig. 6). Thus, this clotting phenomenon is generalizable to different LNP and liposome formulations containing cationic lipids.

### LNPs with cationic lipids aggregate and activate platelets

Having proven that +DOTAP LNPs activate the coagulation cascade, we sought to assess if +DOTAP LNPs also cause activation and aggregation of platelets, since clots can also form due to platelet aggregates^24^. We measured complete blood counts (CBCs) in blood drawn 30 minutes post-injection of +DOTAP LNPs. CBCs showed that +DOTAP LNPs greatly reduced the number of circulating platelets in mice with pre-existing inflammation (nebulized LPS), and also showed a trend towards reduced platelets in naive mice (Fig. 3a and Supplementary Fig. 7). Reduced platelet count, thrombocytopenia, can indicate depletion of platelets from circulation due to incorporation in clots.

**Figure 3:**
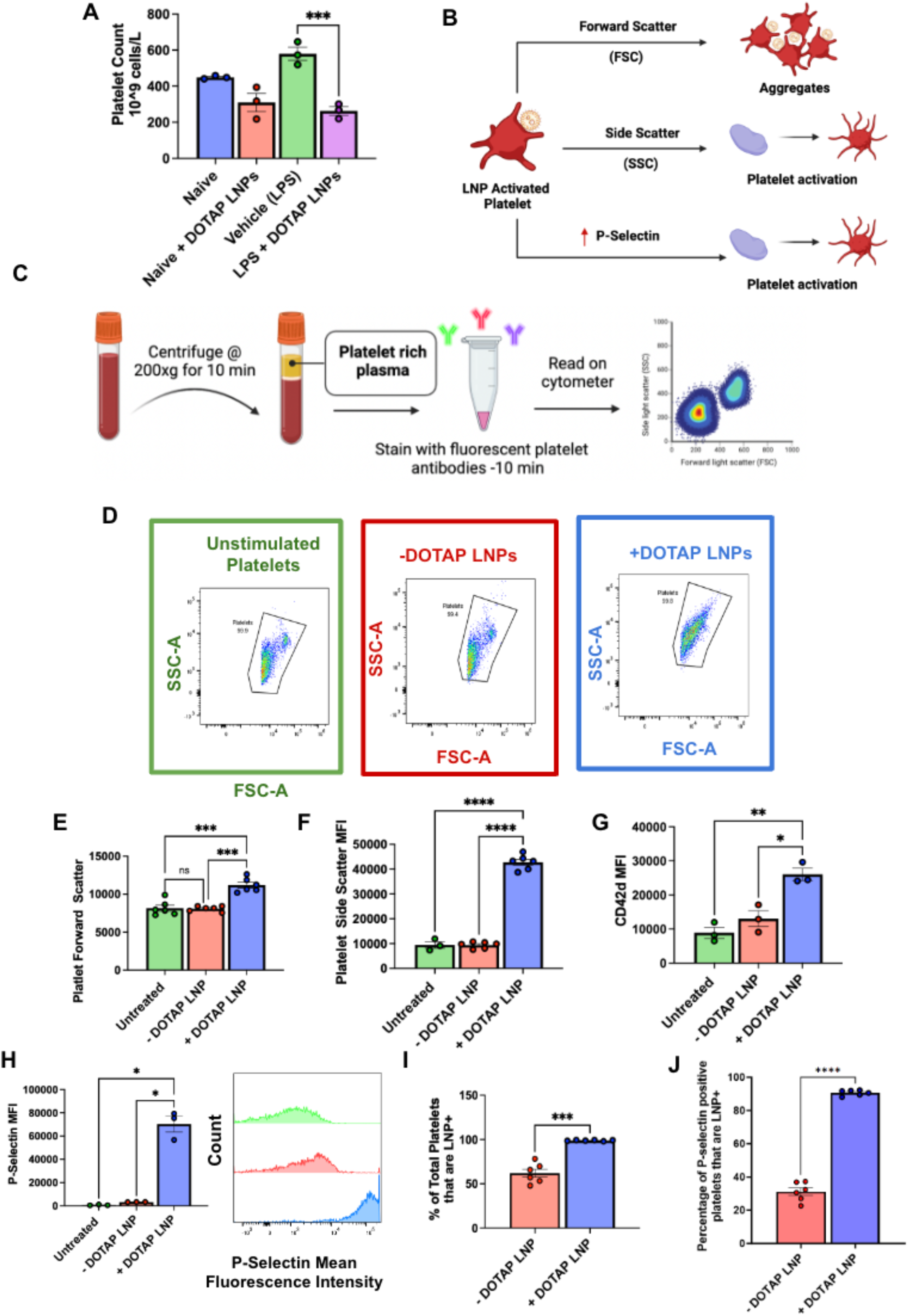
DOTAP LNPs cause platelet activation. (A) +DOTAP LNPs significantly decrease platelet count in naive mice and even more so in mice with pre-existing inflammation (nebulized-LPS). LNPs were IV-injected and platelet count was measured from whole blood 30 minutes later. (B) Schematic depicting what different flow cytometry metrics describe about platelet physiology. (C) Schematic of protocol used for platelet flow cytometry. After isolation of platelet rich plasma (PRP), samples were either untreated, or treated with - DOTAP or +DOTAP LNPs for an incubation period of 30 mins at 37℃. (D) Representative FSC-A vs SSC-A graphs of unstimulated platelets (green) and platelets treated with +DOTAP (blue) or -DOTAP LNPs (red), showing a clear change in scatterplot with +DOTAP LNPs. (E, F) There are significant increases in FSC and SSC upon treatment with +DOTAP LNPs suggesting larger and more complex platelet aggregates (G, H) +DOTAP LNPs cause upregulation of CD42d and P-selectin, based on mean fluorescence intensity (MFI). (I) Platelets were assayed for whether they physically associated LNPs, as measured by the fraction of platelets that were positive for a fluorescent lipid that had been incorporated into the LNPs during synthesis. While 50% of platelets were associated with -DOTAP LNPs, nearly 100% of platelets were physically associated with +DOTAP LNPs. (J) Staining with P-selectin shows that while only 30% of P-selectin positive platelets were associated with - DOTAP LNPs, almost 100% of P-selectin positive platelets were LNP positive as well, showing preference for an activated state. Statistics: n=3-6 and data shown represents mean ± SEM; For I and J, Welch’s t-test was performed ***=p<0.0001. For all other graphs, Brown-Forsythe and Welch’s ANOVA was performed with Dunnett T3 post hoc test. *=p<0.05, ***=p<0.001, ****=p<0.0001

We wanted to investigate if this thrombocytopenia was due to platelet activation by +DOTAP LNPs. To measure platelet activation, we employed flow cytometry. As shown in the schematic of Fig 3b, we used forward scatter (FSC) and side scatter (SSC) measurements to detect larger and more complex platelet aggregates vs. small, isolated platelets. We stained for two specific markers of platelets: glycoprotein V (GPV; CD42d), and P-selectin (CD62p) (Supplementary Fig. 8a, d). CD42d is constitutively found on the surface of all platelets^25^. In resting platelets, P-selectin is generally stored in platelet granules, which are externalized when the platelet becomes activated^25, 26^. Platelet rich plasma (PRP; Fig. 3c) was either left untreated or incubated with fluorescently labeled -DOTAP or +DOTAP LNPs and stained for CD42d and P-selectin.

FSC vs. SSC plots for platelets (CD42d-positive events) in untreated, -DOTAP, and +DOTAP LNP-treated PRP show no difference between untreated and -DOTAP LNPs, but the shape of the +DOTAP LNP graph differs markedly from the other two (Fig. 3d). The mean FSC and SSC values are elevated for platelets in the presence of +DOTAP LNPs, indicating the formation of aggregates (Fig. 3e,f). Further evidence of platelet aggregation induced by +DOTAP LNPs is shown by increased CD42d signal for each CD42d-positive event (Fig. 3g). In the presence of +DOTAP LNPs, each CD42d-positive event is a cluster of platelets, rather than an individual cell, so the mean CD42d signal is increased vs. untreated and -DOTAP LNP samples. Most interesting, however, is the 20-fold increase in P-selectin presentation induced by +DOTAP LNPs (Fig. 3h). Since increased surface presentation of P-selectin is a marker of platelet activation, this demonstrates profound platelet activation induced by +DOTAP LNPs.

Tracing LNP fluorescence in our flow cytometry measurements, we found that 100% of platelets physically associated with fluorescently-labeled +DOTAP LNPs, while only 50% of platelets physically associated with -DOTAP LNPs, showing that the cationic lipid profoundly increases LNP adhesion to platelets (Fig. 3i). Specifically, among platelets that had LNP signal, we assessed P-Selectin signal. For -DOTAP-LNPs, only 30% of the LNP-associated platelets were P-selectin positive, indicating that -DOTAP LNPs had little effect on platelet activation. However, for +DOTAP LNPs, nearly 100% of LNP-positive platelets were also positive for P-selectin, indicating that platelet association with +DOTAP LNPs predicts platelet activation (Fig. 3j).

### DOTAP LNPs bind to fibrinogen, and fibrinogen is required for LNP-induced platelet activation

Having proven that +DOTAP LNPs cause clotting, initiate the coagulation cascade, and activate platelets, we sought to isolate components of the +DOTAP LNP protein corona that might lead to these effects. Upon IV injection, LNPs can form unique protein coronas based on their physicochemical properties and this could alter their functionality^27, 28^. The protein corona can therefore drive the biodistribution and cell-type localization of LNPs. However, the proteins adsorbed on LNPs could also induce side effects: Proteins can undergo conformational changes when adhering to surfaces and aggregates of proteins on surfaces do not behave the same way as proteins in solution, including causing immunogenic or thrombotic side effects^27^.

We hypothesized that +DOTAP LNPs could bind to coagulation proteins in plasma, the most abundant being fibrinogen. Fibrinogen is a soluble protein found in plasma. When it is cleaved by thrombin, fibrinogen undergoes conformational changes and aggregates to form fibrin strands, which in turn form the clot-stabilizing mesh. Fibrinogen/fibrin aggregates on surfaces could cause adhesion and activation of platelets. Fibrinogen has a net negative charge at physiological pH and contains domains with a high concentration of negatively charged residues, such as the E domain^29, 30^. Studies have linked the exposure of certain fibrinogen residues to specific side-effects^31–33^. For example, nanoparticle-induced exposure of a peptide sequence at the C-terminus of the fibrinogen γ-chain interacts with the integrin receptor Mac-1 and induces inflammation^33^. We therefore hypothesized that positively charged DOTAP in LNPs could interact with, and aggregate fibrinogen in a way that could lead to coagulation and platelet activation. This hypothesis is supported by our finding that +DOTAP LNPs decrease the expression of CD41 on platelets (Supplementary Fig. 8b). Since CD41 is the platelet receptor of fibrinogen/fibrin, this indicates that there is a blockage of this marker by fibrinogen induced by +DOTAP LNPs.

We first tested if +DOTAP LNPs bind to fibrinogen in vitro. We added fluorescently labeled fibrinogen and LNPs to plasma anticoagulated with heparin and incubated for 10 minutes. We diluted this sample and performed fluorescence-mode Nanoparticle Tracking Analysis (NTA) to detect nanoparticle-sized aggregates of fluorescent fibrinogen, indicating fibrinogen aggregation on LNPs (Fig. 4a). ∼38-fold more fibrinogen-positive particles were observed in the presence of +DOTAP LNPs vs. -DOTAP LNPs (Fig. 4b-d). This agrees with our findings that +DOTAP LNPs aggregate significantly in plasma and even after incubation with fibrinogen alone (Supplementary Fig. 9 and 10).

**Figure 4:**
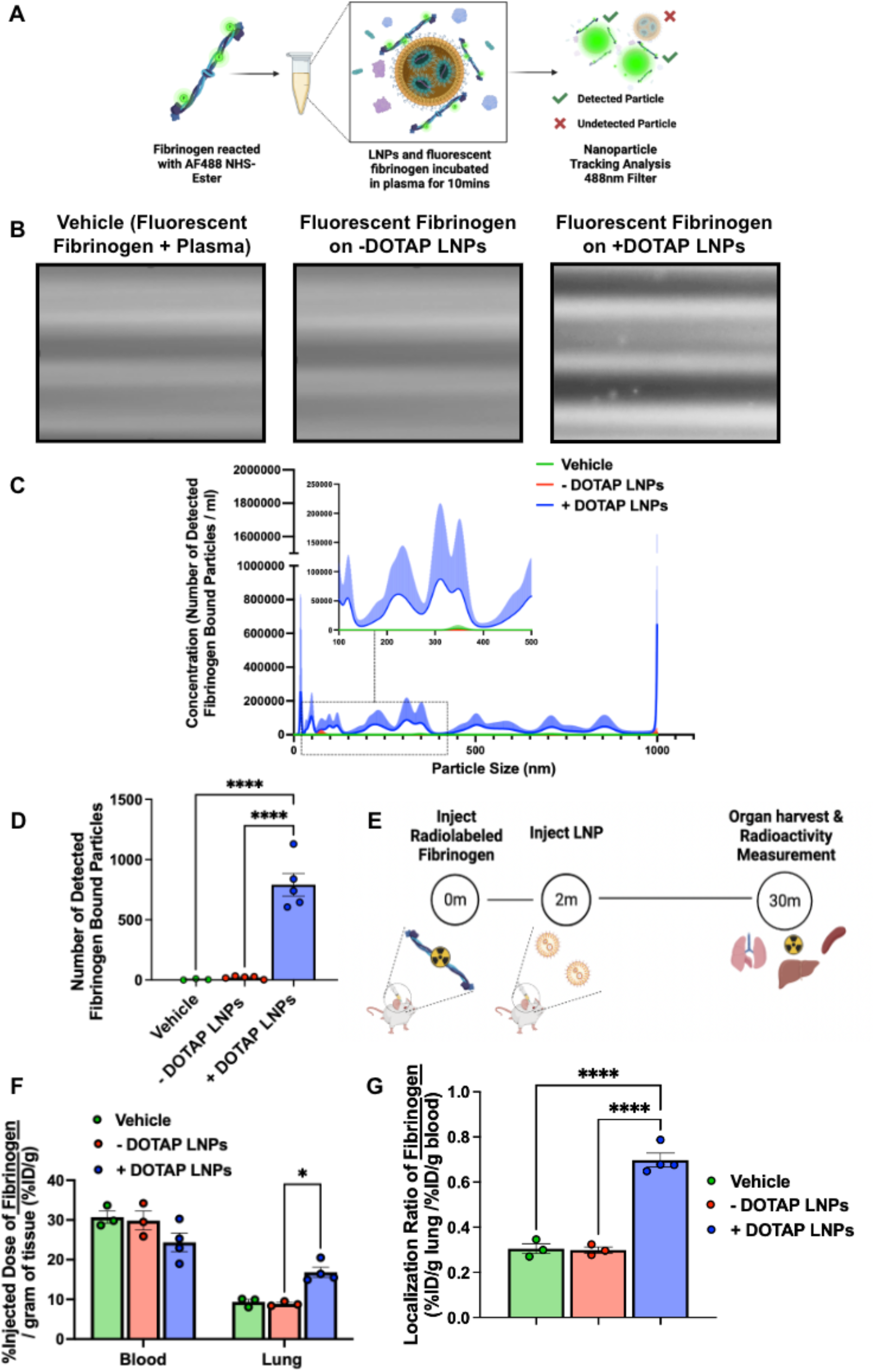
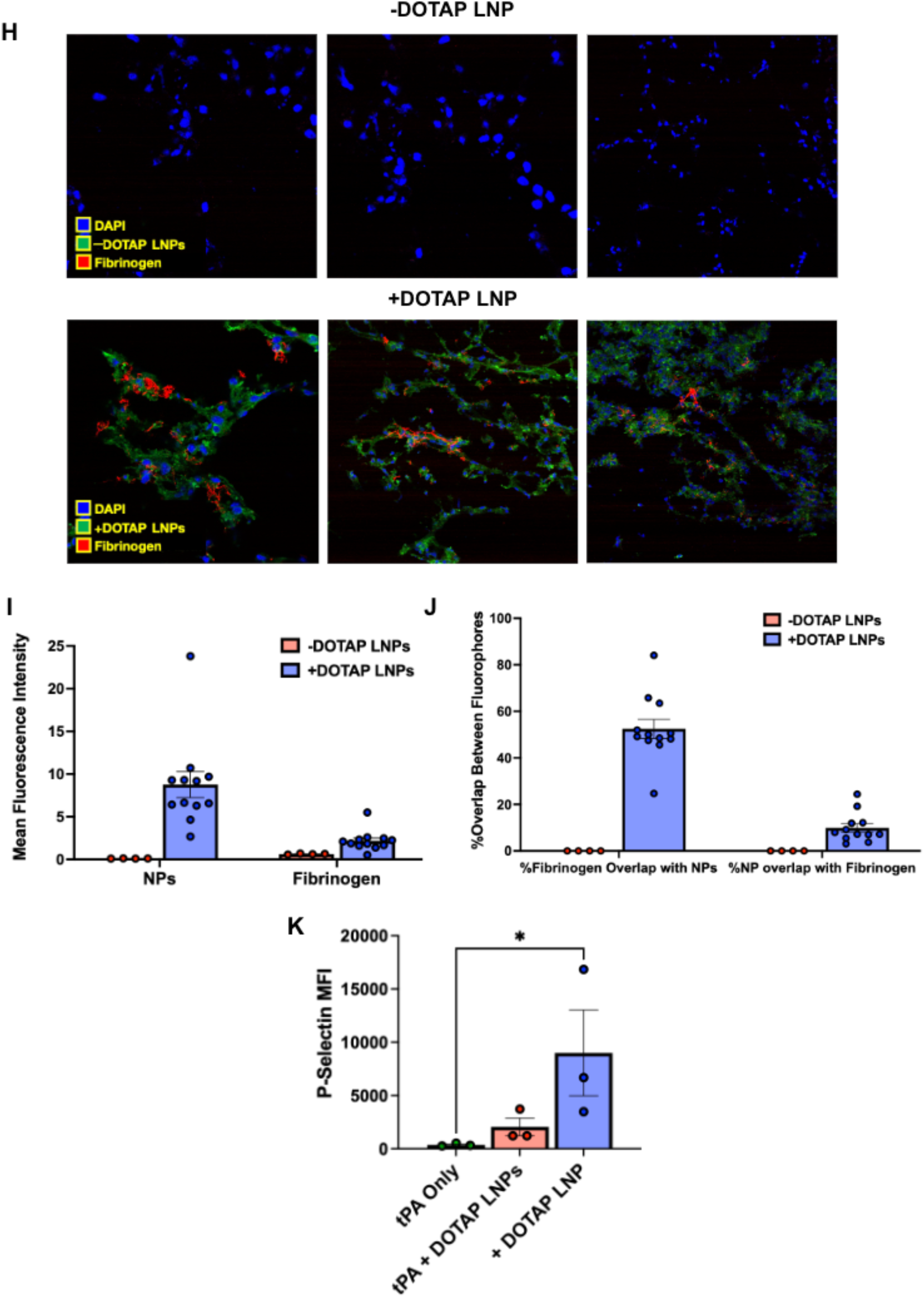
Fibrinogen binds to +DOTAP LNPs, is required for LNP-induced activation of platelets, and is likely the first step in LNP-induced clotting. (A) To measure LNP binding to fibrinogen, we fluorescently labeled fibrinogen, mixed it with plasma and LNPs, and used Nanoparticle Tracking Analysis (NTA) to identify individual nanoparticles that aggregate with fluorescent fibrinogen. (B) Representative NTA images of fluorescent fibrinogen on -DOTAP or +DOTAP LNPs, showing visible binding of fibrinogen to +DOTAP LNPs but no visible binding to -DOTAP LNPs, or in the vehicle control. (C) NTA data was turned into particle size vs concentration histograms, and here we plot fibrinogen-positive particles only. +DOTAP LNPs bind strongly to fluorescent fibrinogen and this leads to LNPs of many different sizes, while -DOTAP LNPs barely have any fibrinogen-positive signal at any particle size. Inset: Size vs concentration histogram ranging from 100nm-500nm. (D) Number of detected fibrinogen bound particles from (C) showing that +DOTAP LNPs generate ∼37.7x more fibrinogen-bound particles than -DOTAP LNPs. (E) Next, we IV-injected radiolabled fibrinogen into mice, followed by LNPs two minutes later. Organs were harvested 30 minutes after LNP injection and the radioactivity in each organ was measured. (F) Amount of radiolabeled fibrinogen in the blood and lungs after the protocol of *E*, shown as the % of injected dose per gram of tissue (%ID/g). +DOTAP LNPs lead to a ∼2x increase in fibrinogen lung uptake, while -DOTAP LNPs do not alter fibrinogen biodistribution compared to control. (G) Localization ratios of fibrinogen from (F) calculated by %ID/g of tissue in the lung divided by that in the blood. +DOTAP LNPs cause a 2.4-fold higher lung localization ratio of fibrinogen compared to -DOTAP LNPs and vehicle. (H) Representative confocal microscopy images of lung sections from mice injected with fluorescent fibrinogen followed by fluorescent LNPs, showing visible fibrinogen aggregates in the lungs of +DOTAP LNP injected mice (n=3 biologically independent animals). (I) Mean fluorescence intensities (MFIs) in the lung sections from (H) of - or +DOTAP LNPs and fibrinogen showing significantly higher MFIs of LNPs and fibrinogen in +DOTAP LNP injected mice compared to -DOTAP LNP mice. (J) From the microscopy images in (H), percentage of fibrinogen signal overlap with that of LNPs or percentage of LNP signal overlap with fibrinogen signal in the presence of - or +DOTAP LNPs shows a strong co-localization of fibrinogen with +DOTAP LNPs. (K) To determine if fibrinogen is necessary for LNP-induced activation of platelets, we added tPA to plasma to deplete the fibrinogen. Fibrinogen depletion nearly completely prevented LNP-induced activation of platelets. Statistics: n=3-4 and data shown represents mean ± SEM; For (F), comparisons between groups were made using 2-way ANOVA with Tukey’s post-hoc test. For all other graphs, comparisons between groups were made using 1-way ANOVA with Tukey’s post-hoc test. *=p<0.05, ***=p<0.001, ****=p<0.0001.

**Figure 5:**
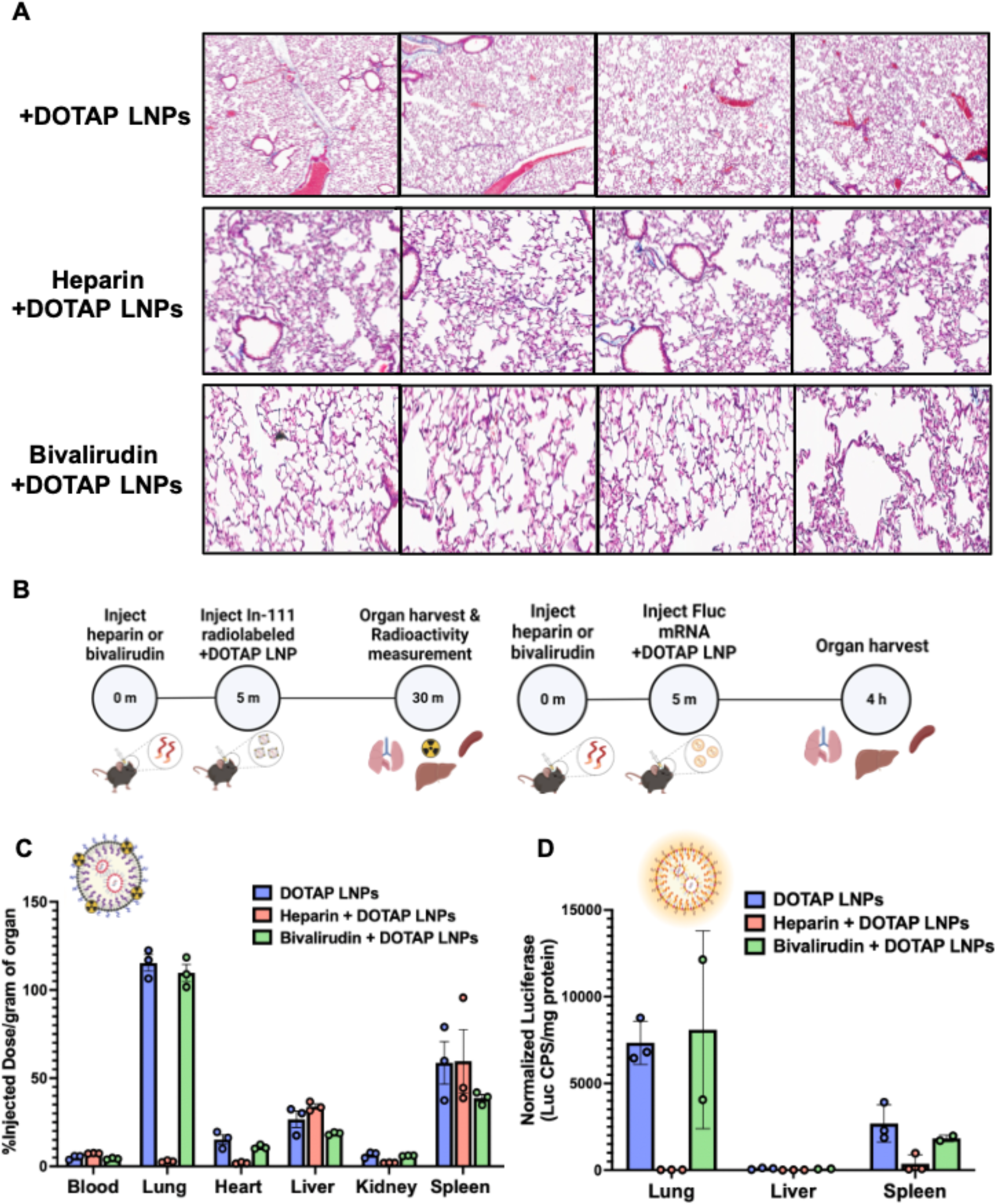
Anticoagulation ameliorates the side-effects of DOTAP LNPs, but only select anticoagulants still allow LNP targeting to the lungs. (A) Histology showing clots in the lungs of mice given +DOTAP LNPs (top row), but not in mice that were given +DOTAP LNPs preceded by heparin (middle row) or bivalirudin (bottom row). (B) Schematics showing protocol for biodistribution (studies using In-111-labeled LNPs (top schematic) and luciferase expression studies. (C) Biodistribution of +DOTAP LNPs with no treatment or pretreated with heparin or bivalirudin. Pretreatment with heparin ablates localization of +DOTAP LNPs to the lung. Interestingly, pretreatment with bivalirudin maintains lung localization. (D) Additionally, bivalirudin further preserves luciferase expression of +DOTAP LNPs whereas heparin pretreatment attenuates this expression capacity. Statistics: n=3 and data shown represents mean ± SEM

We further sought to determine if +DOTAP LNPs bind to fibrinogen in vivo and alter its biodistribution. Fibrinogen was radioactively labeled and injected into mice, followed by + or - DOTAP LNPs two minutes after. Organs were harvested 30 minutes after LNP injection and the radioactivity in each organ was measured (Fig. 4e). While -DOTAP LNPs do not alter fibrinogen biodistribution, there is a ∼2-fold increase in fibrinogen lung uptake in the presence of +DOTAP LNPs (Fig. 4f-g and Supplementary figure 11). A similar +DOTAP LNP-induced increase in lung fibrinogen uptake was observed in mice with pre-existing inflammation (Supplementary Figure 11). Confocal microscopy images of lung sections show visible aggregates of fibrinogen in +DOTAP LNP-injected mice and no visible fibrinogen in -DOTAP LNP injected mice (Fig. 4h). This was quantified across multiple images, with the LNPs and fibrinogen having much more signal in the lungs of +DOTAP vs. -DOTAP LNP mice (Fig. 4i). Furthermore, fibrinogen is strongly associated with +DOTAP LNPs as shown by the percentage of fibrinogen signal overlap with that of LNPs or percentage of LNP signal overlap with fibrinogen signal (Fig. 4j). The data in Figures 4a-j clearly show that fibrinogen binds to +DOTAP LNPs (but not -DOTAP), and this binding correlates with clotting in the lungs.

To further examine the effects of fibrinogen binding to +DOTAP LNPs, we depleted fibrinogen from plasma doped with +DOTAP LNPs. Since fibrinogen depletion would prevent coagulation, we checked if fibrinogen depletion also abrogated +DOTAP LNP-induced platelet activation. PRP was treated with tissue plasminogen activator (tPA) which converts plasminogen to plasmin, causing degradation of fibrinogen. We found that in fibrinogen-depleted plasma, +DOTAP LNPs do not induce significant platelet activation as measured by P-Selectin MFI (Fig. 4k). Thus fibrinogen is *necessary* for +DOTAP LNP-induced platelet activation. This fits with a mechanism in which positively charged lipids first bind to and activate fibrinogen, which then activates platelets, which in turn activate the rest of the coagulation cascade. This fits with prior studies that show that immobilization of fibrinogen on surfaces can expose normally concealed domains that activate platelets^34–37^.

### Anticoagulation ameliorates cationic LNP-induced clotting

Below, we propose and test ways to prevent LNP-induced clotting. To determine whether anticoagulation can reduce the toxicity of +DOTAP LNPs, we compared clot formation between mice treated with +DOTAP LNPs alone and those pre-treated with the clinically used anticoagulants heparin and bivalirudin. Heparin functions by enhancing the activity of endogenous antithrombin while bivalirudin is a direct thrombin inhibitor. Anticoagulants were IV injected 5 minutes prior to +DOTAP LNP injection (62.5U/mouse for heparin and 400ug/mouse for bivalirudin) and LNPs were allowed to circulate for 30 minutes. We found that both heparin and bivalirudin pre-treatment significantly reduce the formation of clots in the lung compared to +DOTAP LNP injection alone, as shown by lung histological samples (Fig. 5a). We then investigated the effect of anticoagulation on +DOTAP LNP biodistribution and mRNA expression. We radiolabeled LNPs with Indium-111 for biodistributions and injected mice with LNPs loaded with luciferase mRNA to trace mRNA expression. Surprisingly, heparin pre-treatment shunted LNP localization from the lung to the liver and spleen and completely attenuated luciferase expression. (Fig. 5b, c). We hypothesize that this unexpected effect is because heparin is a negatively charged polysaccharide, so it could have a charge interaction with +DOTAP LNPs and this could inhibit LNP functionality^38^. However, bivalirudin pre-treatment preserves the localization of +DOTAP LNPs to the lung, as well as the LNP-induced mRNA luciferase expression (Fig 5b,c). These studies illustrate that while the side-effects of +DOTAP LNPs can be ameliorated with anticoagulation, the properties of each anticoagulant must be thoroughly examined to ensure that LNP activity is preserved.

Based on this success with bivalirudin (a direct thrombin inhibitor), we fabricated +DOTAP LNPs with a surface-conjugated direct thrombin inhibitor, PPACK (Supplementary Figure 13a-c)^39^. This might be a more convenient clinical solution than needing to transiently anticoagulant patients with bivalirudin. Clot formation in the lungs was limited in mice treated with PPACK-conjugated +DOTAP LNPs (Supplementary Figure 13d).

### Decreasing the size of DOTAP LNPs prevents fibrinogen binding and clotting

We serendipitously observed that the size of +DOTAP LNPs has a positive correlation with APTT. When we increased LNP size from 100nm to 180nm, APTT increased by ∼1.6-fold. However, when we decreased the LNP size to 80nm, APTT was restored to naive levels indicating no activation of the intrinsic coagulation pathway (Fig. 6a). Similarly, 80nm +DOTAP LNPs did not significantly activate platelets, as measured by P-Selectin MFI (Fig. 6b). This led us to hypothesize that the size of +DOTAP LNPs could affect the amount of fibrinogen molecules that can be adsorbed on the LNP surface. We tested this hypothesis first in vitro using fluorescence NTA to probe for fibrinogen aggregation and found that 80nm +DOTAP LNPs generate ∼79x fewer nanoparticle-associated fibrinogen aggregates than 100nm +DOTAP LNPs (Fig. 6c, d). Additionally, the number of fibrinogen-nanoparticle aggregates detected with 80nm +DOTAP LNPs is not significantly different from that under vehicle conditions, indicating a complete absence of detectable fibrinogen-nanoparticle aggregation. We subsequently tested how 80nm +DOTAP LNPs affect fibrinogen biodistributions in vivo. As previously, we injected mice with radiolabeled fibrinogen followed by LNPs 2 minutes after for a circulation time of 30 minutes. While 100nm +DOTAP LNPs almost double fibrinogen uptake in the lung, 80nm +DOTAP LNPs do not alter fibrinogen biodistribution or localization ratio compared to control (Fig. 6e, f). These data validate our hypothesis that 80nm +DOTAP LNPs do not detectably bind to or cause aggregation of fibrinogen. Since this interaction is associated with thrombosis, these LNPs do not activate platelets or the intrinsic coagulation pathway (Fig. 6g). Furthermore, histological samples from the lungs of LNP-injected mice show that 80nm +DOTAP LNPs do not generate visible clots like 100nm +DOTAP LNPs (Fig. 6h). Finally, 80nm +DOTAP LNPs still preserve their lung-tropism, as shown by their biodistribution and mRNA expression profile (30 minute and 4 hour LNP circulation time respectively) (Fig. 6i, j). These data indicate that by altering the physical properties of +DOTAP LNPs, we can modulate the binding of fibrinogen and the subsequent initiation of clotting, while maintaining lung tropism.

**Figure 6:**
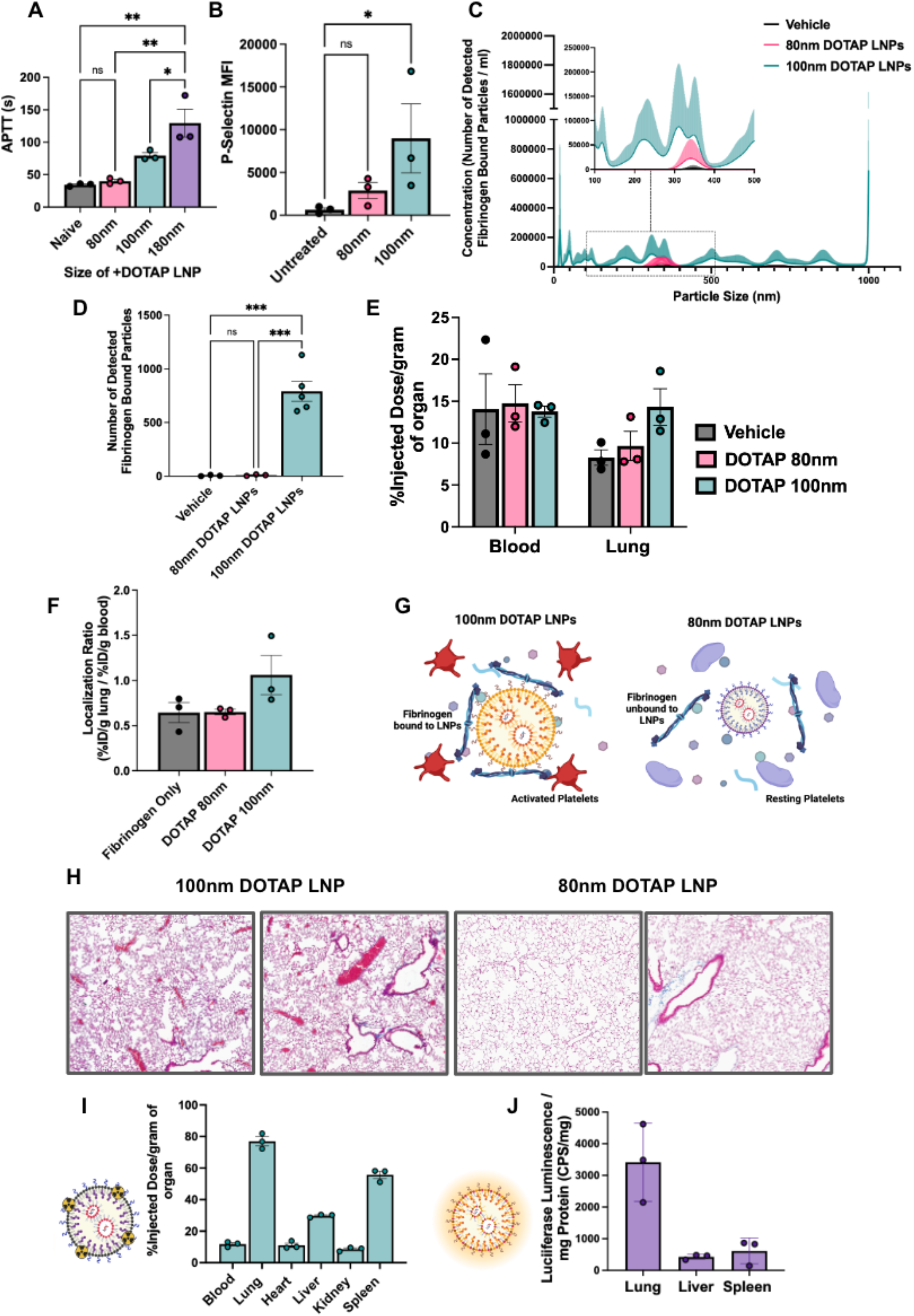
Decreasing the size of DOTAP LNPs prevents fibrinogen binding and the resulting coagulation and platelet activation. (A) +DOTAP LNP size has a positive correlation on coagulation as measured by APTT. 180nm +DOTAP LNPs lead to a ∼1.6-fold increase in APTT compared to standard 100nm +DOTAP LNPs while 80nm +DOTAP LNPs maintain APTT at naive levels. (B) 80nm +DOTAP LNPs do not cause significant platelet activation as measured by P-Selectin MFI. (C) Particle size vs concentration histograms from NTA fluorescent fibrinogen binding assay comparing the number of detected fibrinogen bound particles with 80nm or 100nm +DOTAP LNPs showing that 80nm +DOTAP LNPs bind significantly less fibrinogen compared to 100nm +DOTAP LNPs. Inset: Size vs concentration histogram ranging from 100nm-500nm.(D) Number of detected fibrinogen bound particles from (C) showing that 80nm +DOTAP LNPs generate ∼79x fewer fibrinogen bound particles than 100nm +DOTAP LNPs. The number of fibrinogen bound particles detected with 80nm +DOTAP LNPs is not significantly different from the vehicle. (E) Amount of fibrinogen in the blood and lung with injection of radiolabeled fibrinogen alone (vehicle) or radiolabeled fibrinogen followed by 80nm or 100nm +DOTAP LNPs. 80nm +DOTAP LNPs do not alter fibrinogen biodistribution compared to control. (F) 80nm +DOTAP LNPs do not alter localization ratio of fibrinogen calculated from (E) (%ID/g of tissue in the lung divided by that in the blood). (G) Hypothesized mechanism behind the abrogation of clotting side effects below a size threshold: At a large radius of curvature (≥ 100nm), +DOTAP LNPs present a flat enough surface to bind fibrinogen lengthwise, which then activates platelets. (H) Lung histological samples reveal that 80nm +DOTAP LNPs do not generate visible clots as is the case with 100nm +DOTAP LNPs. 80nm +DOTAP LNPs maintain their biodistribution and lung localization (10ug mRNA/mouse, 30 minutes) (I) and express mRNA primarily in the lung measured by luciferase expression (10ug mRNA/mouse, 4 hours) (J). Statistics: n=3 and data shown represents mean ± SEM. Comparisons between groups were made using 1-way ANOVA with Tukey’s post-hoc test. *=p<0.05, ***=p<0.001, ****=p<0.0001.

## Discussion

Nanoparticles are prone to attack by blood’s defense systems. Two of the major defense systems of blood, complement proteins and immunoglobulins, have been repeatedly shown to attack nanoparticles. But a third blood defense, clotting, is often overlooked by the field of nanomedicine, especially the booming field of LNPs, as this is the first report of LNP-induced thrombosis. Since intravascular clotting leads to potentially deadly acute thrombosis, investigation of clotting side effects should be a standard part of IV nanomedicine development. There have been prior studies showing thrombotic side effects of other types of nanoparticles, but these studies have been largely neglected with the most clinically important nanomedicines today: LNPs. As the field of nanomedicine rapidly moves towards using charged lipids and other physicochemically less-defined mechanisms as a primary tool for targeting, studies of clotting responses to nanomedicines, especially LNPs, are critically timely and necessary.

Among the reasons that the LNP field had not previously observed clotting, a major one is likely the methods used for testing LNPs *in vitro*. *In vitro* studies of blood responses to nanoparticles often use serum, rather than plasma. Plasma is the acellular component of blood. Serum is prepared by depleting clotting factors from plasma, so it is impossible to study clotting responses with serum. For instance, fibrinogen, critical to side effects observed in our studies, is missing from serum. Even in studies of plasma, one misses the critical role of platelets, shown here to serve as a pro-thrombotic surface that is activated by and amplifies the pro-coagulant effects of +DOTAP LNPs. Nanomedicine’s focus on complement as a central source of side effects may also miss upstream coagulation side effects: Thrombin and plasmin activate the complement cascade. *In vitro* and *in vivo* studies that assessed nanoparticle interactions with the acellular and cellular components of blood involved in clotting were necessary to evaluate the deadly side effects identified here, and similar studies may be called for in developing other physicochemical targeting approaches.

Here we have focused on clotting associated with cationic lipids in LNPs, but future studies will need to investigate other nanoparticle properties that induce clotting. In this current study, we showed that LNPs with cationic lipids induce clotting, across a wide range of LNP lipid constituents (varying ionizable lipids, charged lipids, etc). We showed the same clotting side effects in liposomes formulated with cationic lipids. This strongly implicates the presence of cationic moieties as a nanoparticle feature that increases thrombosis risk. But we also identified a nanoparticle feature that reduces thrombosis risk. Varying nanoparticle size had a large effect on clot induction, with a size below ∼100 nm limiting thrombosis induced by +DOTAP LNPs in mice. There are many additional nanoparticle properties that must be tested for their relationship to clotting, including mechanical properties (Young’s modulus, surface fluidity), polymeric brush borders (e.g., PEG), and appended proteins (e.g., for the “old method” of affinity targeting). Such a systematic survey, coupled with biophysics studies of nanoparticle binding to coagulation proteins, will enable the design of safer nanoparticles.

Just as nanoparticle properties must be studied in relation to clotting, we must also investigate how changes in blood itself might affect nanoparticle-associated clotting. While these studies were done in mice, we must now assay for nanoparticle-associated clotting in human blood, as well as the large animals commonly used in preclinical efficacy and toxicity studies (pig, sheep, and non-human primates), noting that small animal models often do not capture fatal embolic side effects that emerge in larger animals and humans. We should also investigate how various disease states and medications that affect the blood might predispose or protect from nanoparticle-associated clotting. Here we showed that pre-existing inflammation worsens the magnitude of LNP-induced lung injury, but more in-depth mechanistic studies are required, as well as testing in other relevant disease states, such as common procoagulant genotypes (Factor V Leiden, Proteins C & S deficiency), cancers with clotting predispositions (noting that clotting is associated with metastasis), and other inflammatory diseases. Indeed, fibrinogen is an “acute-phase reactant,” meaning that its concentration in plasma goes up dramatically during inflammation (and many cancers), thus making it more likely to bind to LNPs. These studies in different disease states are essential, because many of the proposed applications of LNPs are for cancer and inflammatory diseases, and thus nanoparticle-induced clotting could become more prominent and dangerous, perhaps occurring even for smaller LNPs or LNPs without positively charged lipids.

Finally, more mechanistic studies are needed to further understand nanoparticle-induced clotting. Here we showed that fibrinogen binds to positively charged LNPs, and that this binding leads to downstream thrombosis steps (Fig. 4). We hypothesize that fibrinogen binding to LNPs changes its conformation to resemble its activated form. Such surface-bound fibrinogen/fibrin is known to bind to GPIIB/IIIA on platelets, immobilizing and leading to activation of the platelets. Indeed, we showed that fibrinogen-binding +DOTAP LNPs induce platelet aggregation and activation (Fig. 3). But many mechanistic questions remain, such as why fibrinogen binding is so sensitive to the LNP radius of curvature, and whether clinically-used platelet inhibitors (aspirin, clopidogrel, and especially the GPIIB/IIIA inhibitor tirofiban) would prevent LNP-induced clotting. Such studies could improve the techniques used to prevent LNP-associated clotting.

## Methods

### Materials

DOTAP (1,2-dioleoyl-3-trimethylammonium-propane (chloride salt)), DOPE (1,2-dioleoyl-*sn*-glycero-3-phosphoethanolamine), cholesterol, DMG-PEG 2000 (1,2-dimyristoyl-rac-glycero-3-methoxypolyethylene glycol-2000), DPPC (dipalmitoyl phosphatidylcholine), 18:1 PE TopFluor AF 594 (1,2-dioleoyl-sn-glycero-3-phosphoethanolamine-N-(TopFluor® AF594) (ammonium salt)), 18:0 PE-DTPA (1,2-distearoyl-sn-glycero-3-phosphoethanolamine-N-diethylenetriaminepentaacetic acid (ammonium salt)), and DSPE-PEG2000-azide (1,2-distearoyl-sn-glycero-3-phosphoethanolamine-N-[azido(polyethylene glycol)-2000] (ammonium salt)) were purchased from Avanti Polar Lipids. Ionizable lipids cKK-E12, SM-102, ALC-0315, and C12-200 were purchased from Echelon Biosciences. Anti-mouse BV421 CD41, PE-Cy7 CD62P AF700 Ly6G and APC CD42d were purchased from Biolegend. Indium-111 Chloride (In-111) was purchased from BWXT Medical. A modified Lowry assay kit (DC Protein Assay) was purchased from Bio-Rad Laboratories. tPA was purchased from Millipore.

### Animals

All animal studies were carried out in accordance with the Guide for the Care and Use of Laboratory Animals (National Institutes of Health, Bethesda, MD), and all animal protocols were approved by the University of Pennsylvania Institutional Animal Care and Use Committee. All animal experiments were carried out using male, 6–8 week old C57BL/6 mice (20–25 g) (The Jackson Laboratory, Bar Harbor, ME). The mice were maintained at 22-26°C and adhered to a 12/12h dark/light cycle with food and water ad libitum.

### Nanoparticle Formulation

LNPs were formulated using the microfluidic mixing method. An organic phase containing a mixture of lipids dissolved in ethanol at a designated molar ratio (Supplementary Table 1) was mixed with an aqueous phase (50 mM citrate buffer, pH 4) containing Luciferase mRNA that was either purchased by TriLink (most experiments) or made in-house via in vitro transcription (IVT)^40^ at a flow rate ratio of 1:3 and at a total lipid/mRNA weight ratio of 40:1 in a microfluidic mixing device (NanoAssemblr Ignite, Precision Nanosystems). LNPs were dialysed against 1× PBS in a 10 kDa molecular weight cut-off cassette for 2 h, sterilized through a 0.22 μm filter and stored at 4 °C.

To manufacture LNPs of 180 nm, we varied the flow rate of the NanoAssemblr Ignite, from 6 mL/min (for 100 nm) to 1 mL/min (180nm). We were unable to produce uniform LNPs < 100 nm with TriLink mRNA, but with in-house IVT mRNA, we made LNPs of 80 nm at a flow-rate of 6ml/min.

Liposomes were synthesized using the thin film hydration method. Lipids were dissolved in chloroform and combined in a borosilicate glass tube. Chloroform and ethanol were evaporated by blowing nitrogen over the solution until visibly dry (∼15 minutes) then putting the tube under vacuum for >1hr. Dried lipid films were hydrated with 1X PBS, pH 7.4 to a total lipid concentration of 20mM. The rehydrated lipid solution was vortexed and sonicated in a bath sonicator until visually homogeneous (approximately 1 minute each of vortexing and sonication). The solution was then extruded twenty-one times through a 0.2 μm polycarbonate filter.

### Nanoparticle Characterization

Dynamic light scattering measurements of hydrodynamic nanoparticle size, distribution, polydispersity index, and zeta potential were made using a Zetasizer Pro ZS (Malvern Panalytical). LNP RNA encapsulation efficiencies and concentrations were measured using a Quant-iT RiboGreen RNA assay (Invitrogen).

### Nebulized LPS Model

Mice were exposed to nebulized LPS in a ‘whole-body’ exposure chamber, with separate compartments for each mouse (MPC-3 AERO; Braintree Scientific, Inc.; Braintree MA). To maintain adequate hydration, mice were injected with 1mL of sterile saline, 37C, intraperitoneally, immediately before exposure to LPS. LPS (L2630-100mg, Sigma Aldrich) was reconstituted in PBS to 10mg/mL and stored at −80C until use. Immediately before nebulization, LPS was thawed and diluted to 5mg/mL with PBS. LPS was aerosolized via mesh nebulizer (Aerogen, Kent Scientific) connected to the exposure chamber (NEB-MED H, Braintree Scientific, Inc.). 5mL of 5mg/mL LPS was used to induce the injury. Nebulization was performed until all liquid was nebulized (∼20 minutes).

### TAT and Blood Count Measurements

LNPs were injected into naïve or nebulized-LPS injured mice for a circulation time of 30 minutes. Blood was collected from mice into tubes containing EDTA. Blood cells were analyzed using an Abaxis VetScan HM5 Hematology Analyzer for complete blood counts. Blood was then centrifuged at 1500 x g for 10 minutes and plasma was collected and analyzed with a TAT ELISA kit (Abcam) according to the manufacturer’s instructions.

### Lung Histology

LNPs were injected into naïve mice for a circulation time of 30 minutes. After exsanguination and perfusion via the right ventricle with ≈5 mL of phosphate-buffered saline (PBS) at a constant pressure of 25 cm H2O, whole lungs were inflated and fixed with neutral buffered 10% formalin. Paraffin-embedded 5 μm lung sections were stained and imaged with Masson’s Trichrome by the Pathology Core Laboratory of Children’s Hospital of Philadelphia.

### In vitro PT and APTT assays

PT and APTT measurements were carried out with Pacific Hemostasis instruments and reagents at a temperature of 37℃ and time to clot measurements are automatically recorded based on the attenuation of an oscillating stainless steel ball placed in the sample by the formation of a clot. Blood was first collected from naïve mice and anticoagulated with 3.2% sodium citrate at a ratio of 10:1, blood:citrate, vol:vol. Blood sample was then centrifuged at 1500 x g for 10 minutes and the plasma was collected. 50ul of plasma was incubated with 0.5ug of mRNA in LNPs for 5 minutes. For the PT assay, 50ul of LNP-treated plasma was diluted up to 100ul with PBS and 200ul of PT reagent (Pacific Hemostasis) was added and the time to clot was automatically measured. For the APTT assay, 50ul of LNP-treated plasma was diluted up to 100ul of PBS and 100ul of APTT-XL reagent (Pacific Hemostasis) was added for an incubation time of 4 minutes. 100ul of 0.02M calcium chloride was then added and the time to clot was automatically measured.

### Nanoparticle Tracking Analysis Assay for Fibrinogen Aggregation with LNPs

To prepare fluorophore-labeled fibrinogen, mouse fibrinogen (Innovative Research) was incubated with NHS ester Alexa Fluor 488 (ThermoFisher) at 1:10 mol:mol ratio in PBS at 4℃ for 16 hours. Afterwards, excess fluorophore was removed from fibrinogen by 3-fold passage against a 10 kDa molecular weight cutoff centrifugal filter (Amicon) with PBS washing between passages. After fibrinogen recovery from the centrifugal filter, spectrophotometer measurement of optical density at 280 nm (Nanodrop) determined fluorescent fibrinogen concentration and optical density measurement at 488 nm determined the number of fluorophores per fibrinogen.

Immediately before experiments, LNP concentrations were determined by nanoparticle tracking analysis (Nanosight, Malvern). In a total reaction volume of 40 μL, 4E10 LNPs were combined with 20 μL of heparinized mouse serum (a pooled sample obtained from n=3 mice) and fluorescent fibrinogen was doped into the solution at a final physiologically relevant concentration of 3 mg/mL. Fluorescent fibrinogen, serum, and LNPs were incubated in the dark at room temperature for 10 minutes. Fluorescent fibrinogen was also added to serum solutions at identical concentration, without LNPs, verifying that the fluorescent fibrinogen did not detectably adhere to endogenous serum components. The fibrinogen-serum-LNP reactions were terminated by 1:250 dilution in PBS and the diluted suspensions were used for nanoparticle tracking analysis. Nanoparticle tracking analysis was conducted with a 488 nm excitation laser and a 500 nm long pass filter to image and track Alexa Fluor 488 signal from fluorescent fibrinogen on LNPs. Automated analysis of fluorescence nanoparticle tracking data in Malvern Nanosight software used a uniform detection threshold of 5 for all samples. For both fluorescence data and scattering data, three to five technical replicates were obtained for each sample and an average of those replicates was taken as representative of the size-concentration profile for each sample.

### Radiolabeling and Biodistributions

For biodistribution studies, nanoparticles were traced with In-111 as previously described^41^. Nanoparticles were produced as described above with 0.1 mol% of 18:0 PE-DTPA (a chelator-containing lipid) using metal free buffers. Trace metals were removed from the buffers using a Chelex 100 resin, per manufacturer’s instructions, to prevent unwanted occupancy of the chelator. In-111 chloride was added to the nanocarriers at a specific activity of 1 uCi of In-111 per 1 umol of lipid. The mixture was incubated at room temperature for 30 minutes. Then, unincorporated In-111 was removed using a Zeba Spin desalting column. The removal of unincorporated In-111 was verified using thin film chromatography (TLC). A 1 uL sample of nanoparticles was applied to the stationary phase (silica gel strip). The strip was placed in the mobile phase of 10mM EDTA until the solvent front was 1 cm from the end of the strip (∼10 minutes). The strip was cut 1 cm above the initial sample location. In-111 chelated to the nanoparticles stays at the origin, while unchelated In-111 travels with the solvent front. The activity in each section was measured using a gamma counter. The percent of In-111 chelated to the nanoparticles was calculated as the activity in the origin strip divided by the total activity in both strips. For all experiments, >95% of In-111 was chelated to the nanoparticles.

For biodistributions of fibrinogen, the protein was radiolabeled with Iodine-125 using the Iodogen method. Tubes coated with 100 μg of Iodogen reagent were incubated with fibrinogen (2 mg/mL) and Na125I (115μCi/μg protein) for 5 min on ice. Unincorporated I-125 was removed with a Zeba column followed by TLC analysis as described above.

Nanoparticle biodistributions, In-111 labeled nanoparticles were injected into mice for a circulation time of 30 minutes. For fibrinogen biodistributions, I-125 fibrinogen was allowed to circulate for 2 minutes before the injection of nanoparticles for 30 minutes before harvest. The radioactivity in each organ was then read with a gamma counter (Wizard2, PerkinElmer).

### Confocal Microscopy

Healthy mice were injected with 150µg of fibrinogen fluorescently labeled with AF647-NHS-Ester. Two minutes after, mice were injected with fluorescent + or -DOTAP LNPs formulated with 0.3 mol % of 18:1 PE TopFluor AF 594. After a 30-minute circulation the animals were sacrificed and perfused with 5ml of cold PBS and lungs were harvested and freshly frozen. 10 µm lung slices were cryosectioned for imaging using Leica TCS SP8 confocal microscopy.

### Luciferase Delivery

Luciferase mRNA LNPs fabricated as described above were injected into mice for a circulation time of 4 hours. Select organs were then flash frozen until the day of analysis or homogenized immediately. Samples were suspended in 900 uL of homogenization buffer (5mM EDTA, 10mM EDTA, 1:100 diluted stock protease inhibitor (Sigma), and 1x (PBS), samples were then loaded with a steel bead (Qiagen), then placed in a tissue homogenizer (Powerlyzer 24, Qiagen) using the following settings: Speed (S) 2000 rpm, 2 Cycles (C), T time 45 sec, and pause for 30 sec). After this, 100 uL of lysis buffer (10% Triton-X 100 and PBS) was added into each tube and then allowed to incubate for 1 hr at 4C. After this, they were immediately transferred into fresh tubes, and sonicated, using a point sonicator to remove in excess DNA, using an amplitude of 30%, 5 cycles of 3 secs on/off. After this, samples were then centrifuged at 160,000 x g for 10 minutes. The resultant lysate is either frozen or prepared for luminometry analysis.

For luciferase expression 20 uL of undiluted sample was loaded onto a black 96 well-plate then 100uL luciferin solution (Promega) added immediately before reading on a luminometer (Wallac). Last, a Lowry assay (Bio-Rad) is performed according to manufacturer specification using diluted samples, specifically a 1:40 dilution for lung and spleen tissues and a 1:80 dilution for liver tissues. Final luminescence readings were then normalized based on total protein concentration obtained from Lowry Assay

### Platelet Flow Cytometry

Platelet rich plasma (PRP) was collected by first adding 15 uL of 10 mg/mL bivalirudin as well as flushing 23G collection needles with the same solution bivalirudin, collecting between 500-1000 uL of blood per mouse. After blood draw, samples were then centrifuged at room temperature (RT) at 200 xg for 10 minutes. After centrifugation, PRP was collected, taking care to not disturb the resultant pellet. After collection, samples are immediately read on a hematology analyzer (Vetscan HM2, Abaxis) to obtain platelet count. Resultant platelets were then aliquoted to achieve a total of 400,000 platelets per analysis with Tyrode’s buffer. These platelet samples were then incubated with 0.5ug mRNA dose of AlexFluor 594 labeled -DOTAP or +DOTAP LNPs for 30 minutes at 37. After incubation with nanoparticles, samples were stained with flow cytometry antibodies against CD41, CD42d, and p-selectin and allowed to incubate at RT for 10 minutes. Samples were then immediately diluted with 350 uL of Tyrode’s buffer and read on LSR Fortessa (BD Bioscience). Analysis was then completed using FloJo and gating strategy employed is found in Supplementary Fig. 8. For fibrinogen depleted plasma, 0.278 mg of tPA was added per mL of platelet free plasma, followed by 3h incubation at 37°C.

### Statistics

All results are expressed as mean ± SEM unless specified otherwise. Statistical analyses were performed using GraphPad Prism 8 (GraphPad Software) * denotes p<0.05, ** denotes p<0.01, *** denotes p<0.001, **** denotes p<0.0001.

## Acknowledgements

Research reported in this publication was supported by the American Heart Association under Grant 23PRE1014444 (to S.O), Ruth L. Kirschstein National Research Service Award (NRSA) F31HL154662 (to M.E.Z) American Heart Association under Grant 916172 (to J.N.), and Grant NIH R01 HL157189 (to V.M, J.W.M, and J.S.B).

The data for this manuscript were generated in the Penn Cytomics and Cell Sorting Shared Resource Laboratory at the University of Pennsylvania and is partially supported by the Abramson Cancer Center NCI Grant (P30 016520). The research identifier number is RRid:SCR_022376

---

**Supplementary Fig. 1:**
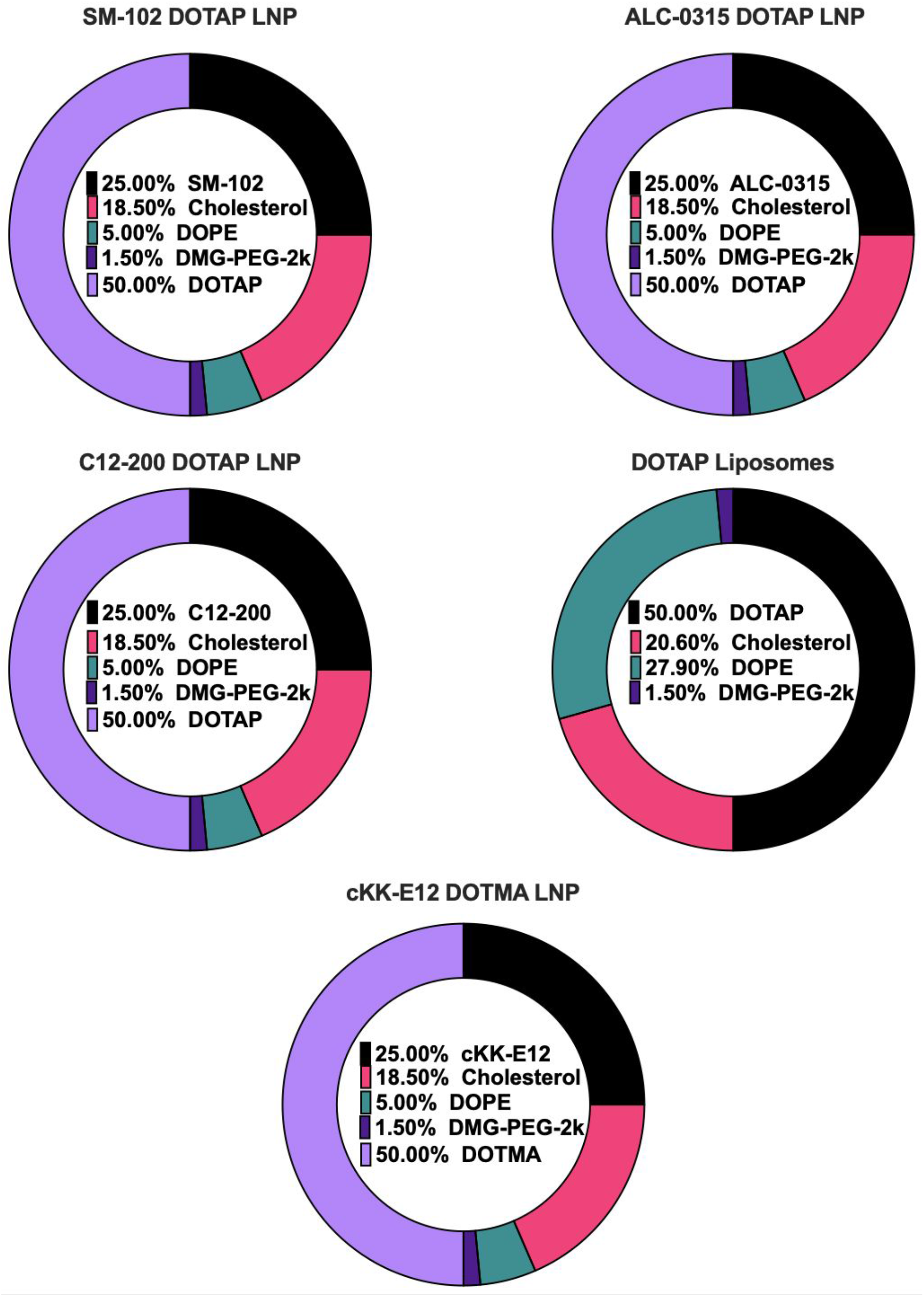
Lipid formulations and molar percentages of DOTAP LNPs fabricated with the ionizable lipids SM-102, ALC-0315, and C12-200 as well as DOTAP liposomes and DOTMA LNPs.

**Supplementary Table. 1:** Size, PDI and zeta potential measurements of nanoparticle formulations obtained with dynamic light scattering.

| Formulation | Size (nm) | PDI | Zeta Potential (mV) |
| --- | --- | --- | --- |
| SM-102 DOTAP LNP | 85.24 | 0.1101 | 9.43 |
| ALC-0315 DOTAP LNP | 157.9 | 0.08745 | 15.19 |
| C12-200 DOTAP LNP | 106.1 | 0.1963 | 12.02 |
| DOTAP Liposomes | 108.9 | 0.2078 | 10.86 |
| cKK-E12 DOTMA LNP | 121.7 | 0.1359 | 5.85 |
| cKK-E12 DOTAP LNP (Smaller) | 79.23 | 0.1258 | 13.94 |
| cKK-E12 DOTAP LNP (Larger) | 179.9 | 0.07139 | 13.69 |

**Supplementary Fig. 2:**
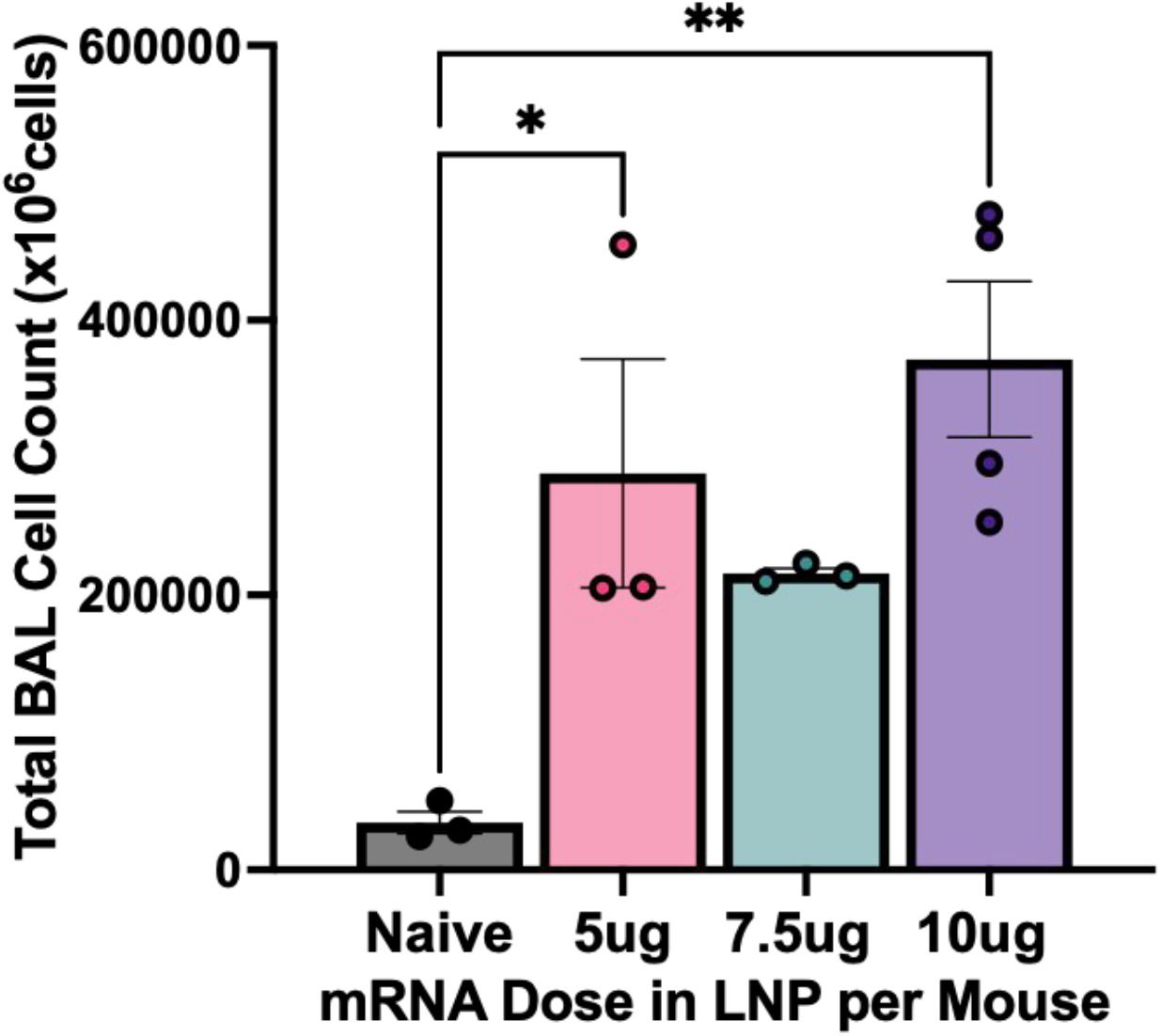
Dose-response of the effect of +DOTAP LNP dose on cell count in bronchoalveolar lavage (BAL) fluid in naïve mice shows a dose-dependent increase.

**Supplementary Fig. 3:**
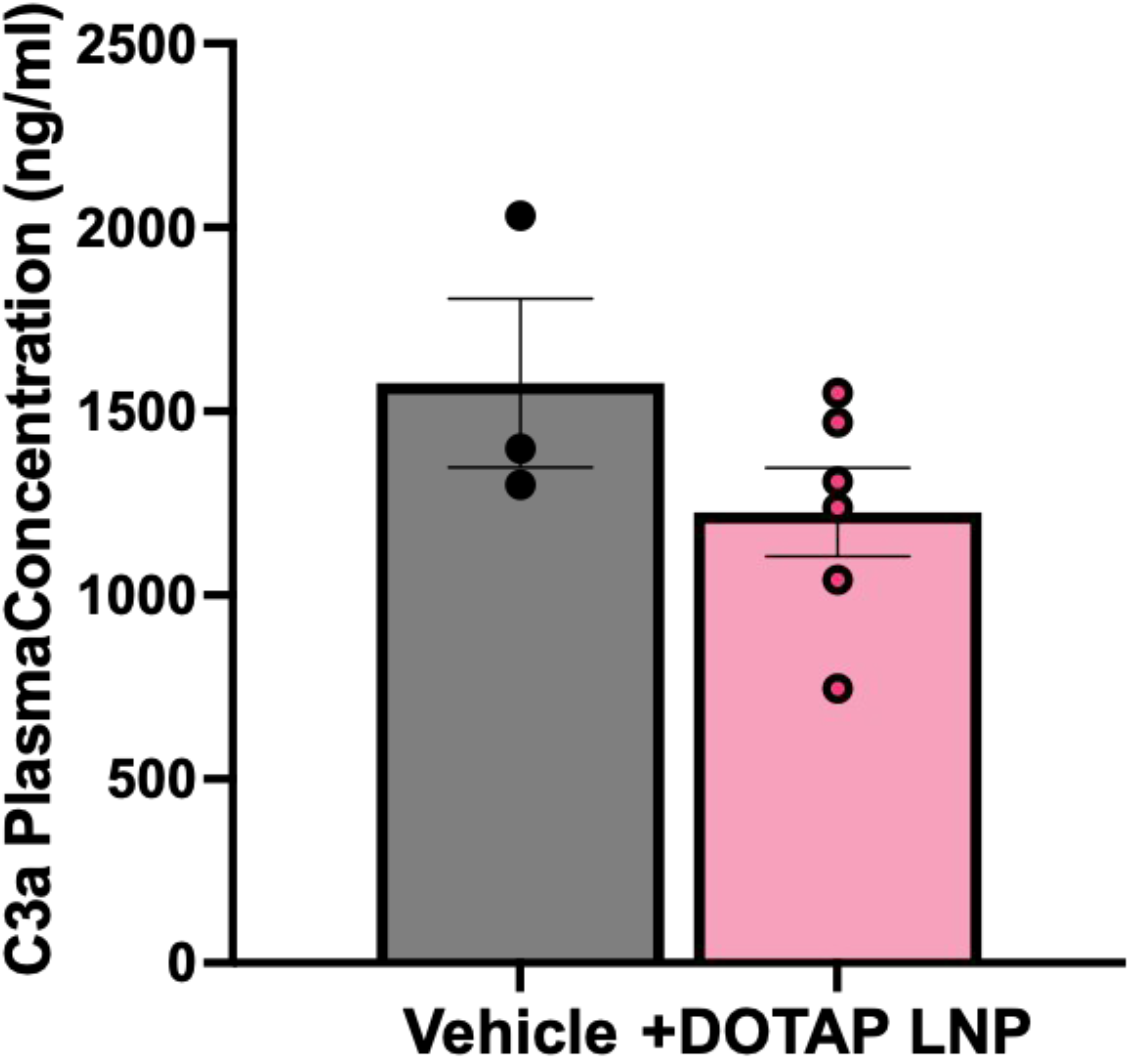
+DOTAP LNPs do not increase the plasma concentration of the complement protein C3a 30 minutes after IV injection into naive mice. C3a concentration was measured by conducting a Mouse C3a ELISA (BD Pharmigen).

**Supplementary Fig. 4:**
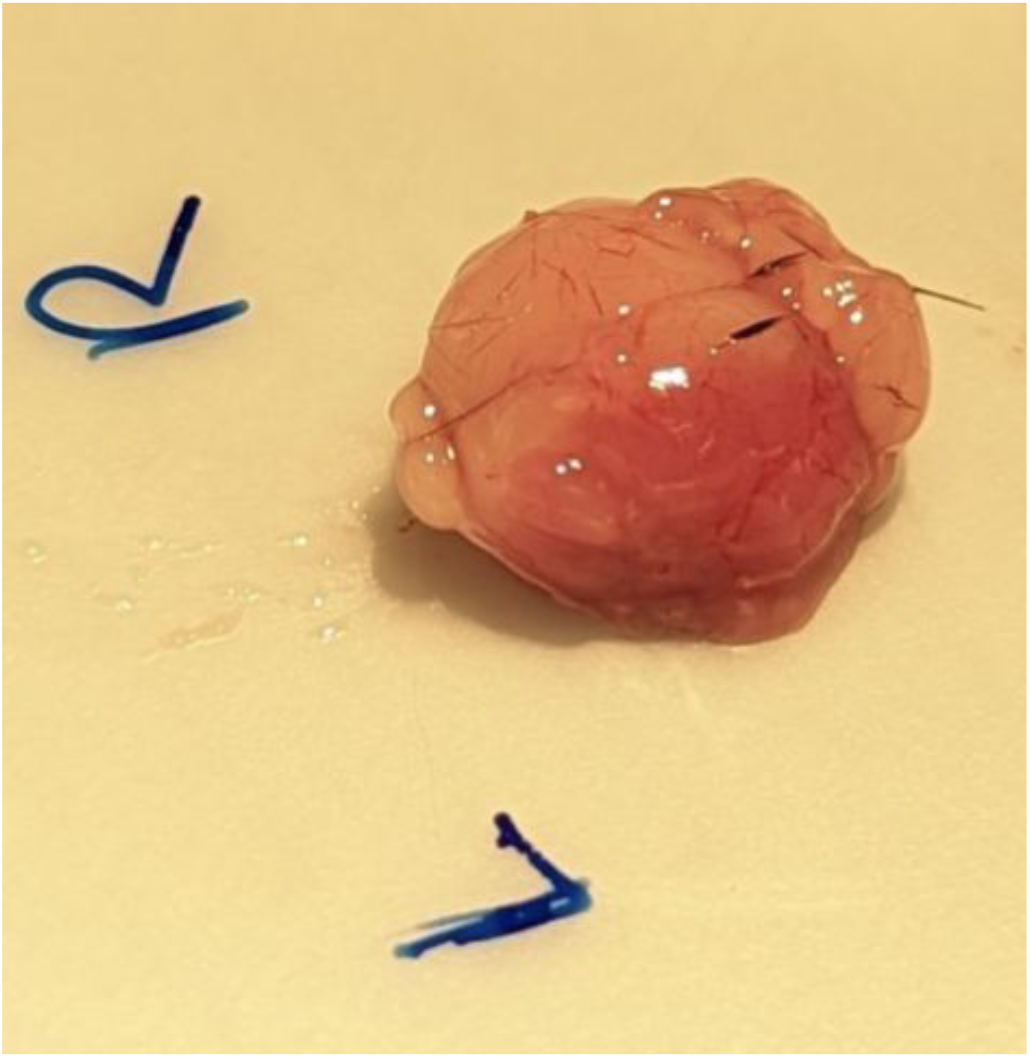
+DOTAP LNPs cause visible indications of clotting in the brain when injected through the carotid artery for a circulation time of 30 minutes.

**Supplementary Fig. 5:**
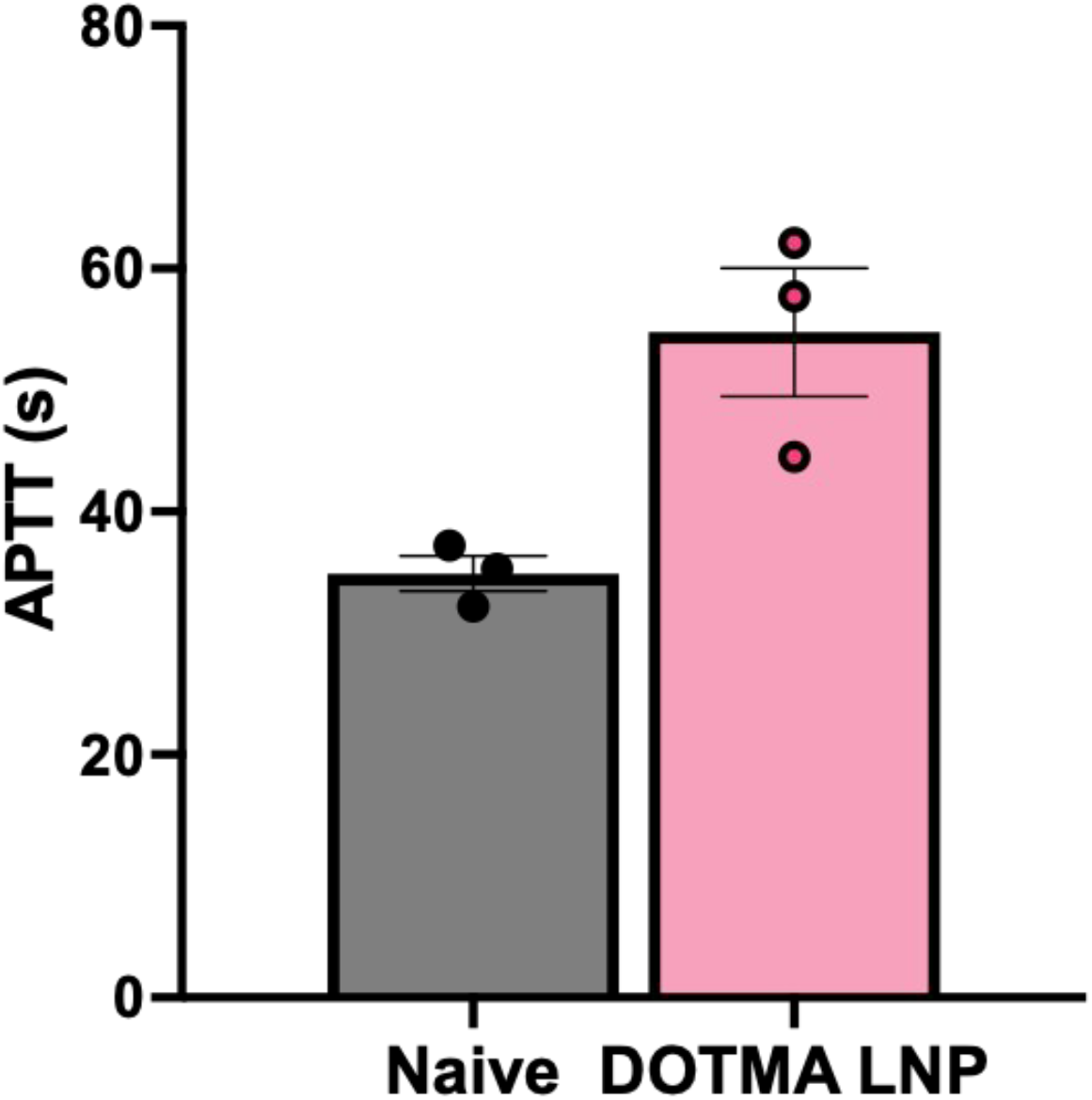
LNPs formulated with the cationic lipid DOTMA also increase APTT *in vitro*.

**Supplementary Fig. 6:**
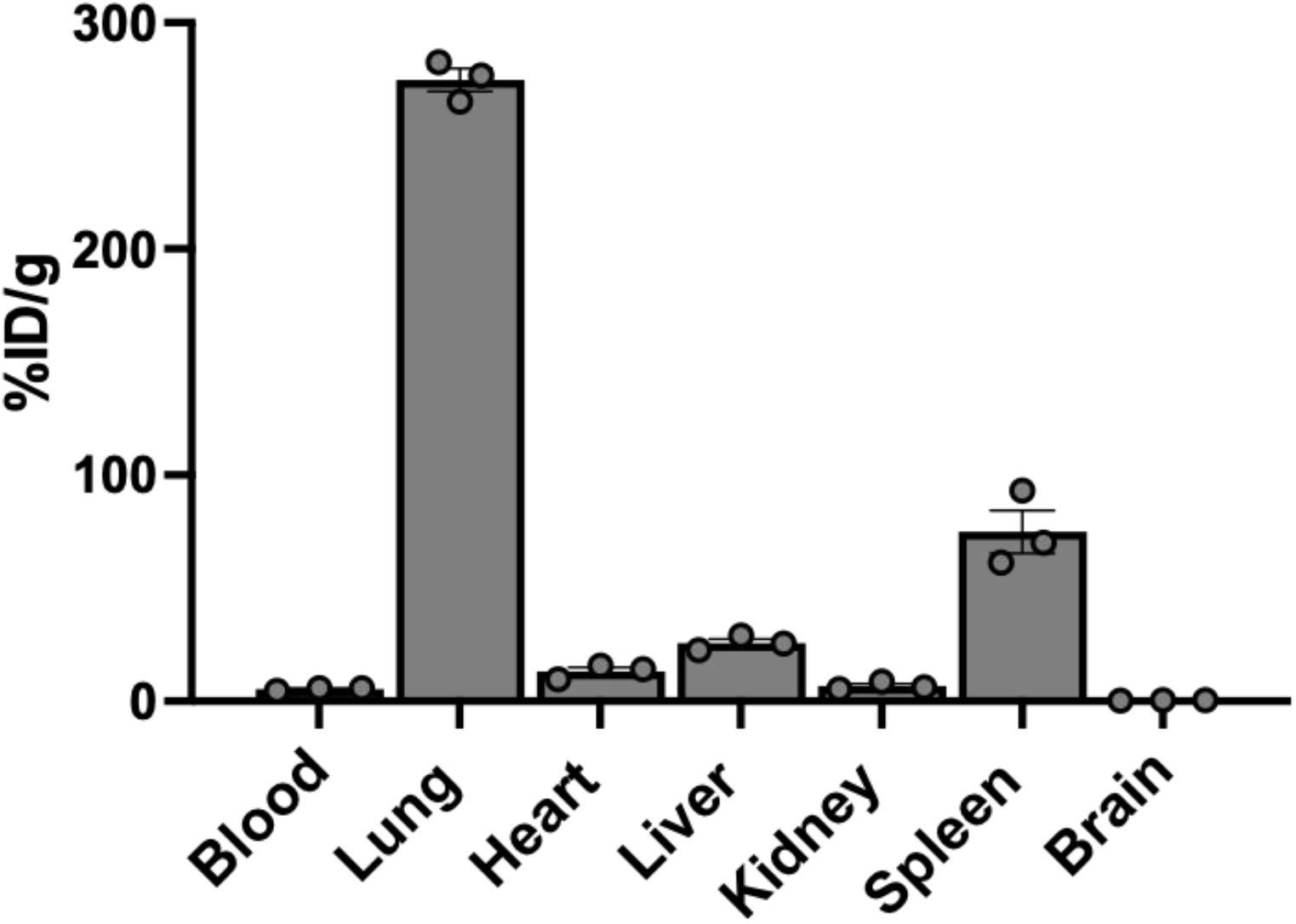
DOTAP liposomes also localize primarily to the lungs when injected into naïve mice. DOTAP liposomes were radiolabeled and intravenously injected for a circulation period of 30 minutes.

**Supplementary Fig. 7:**
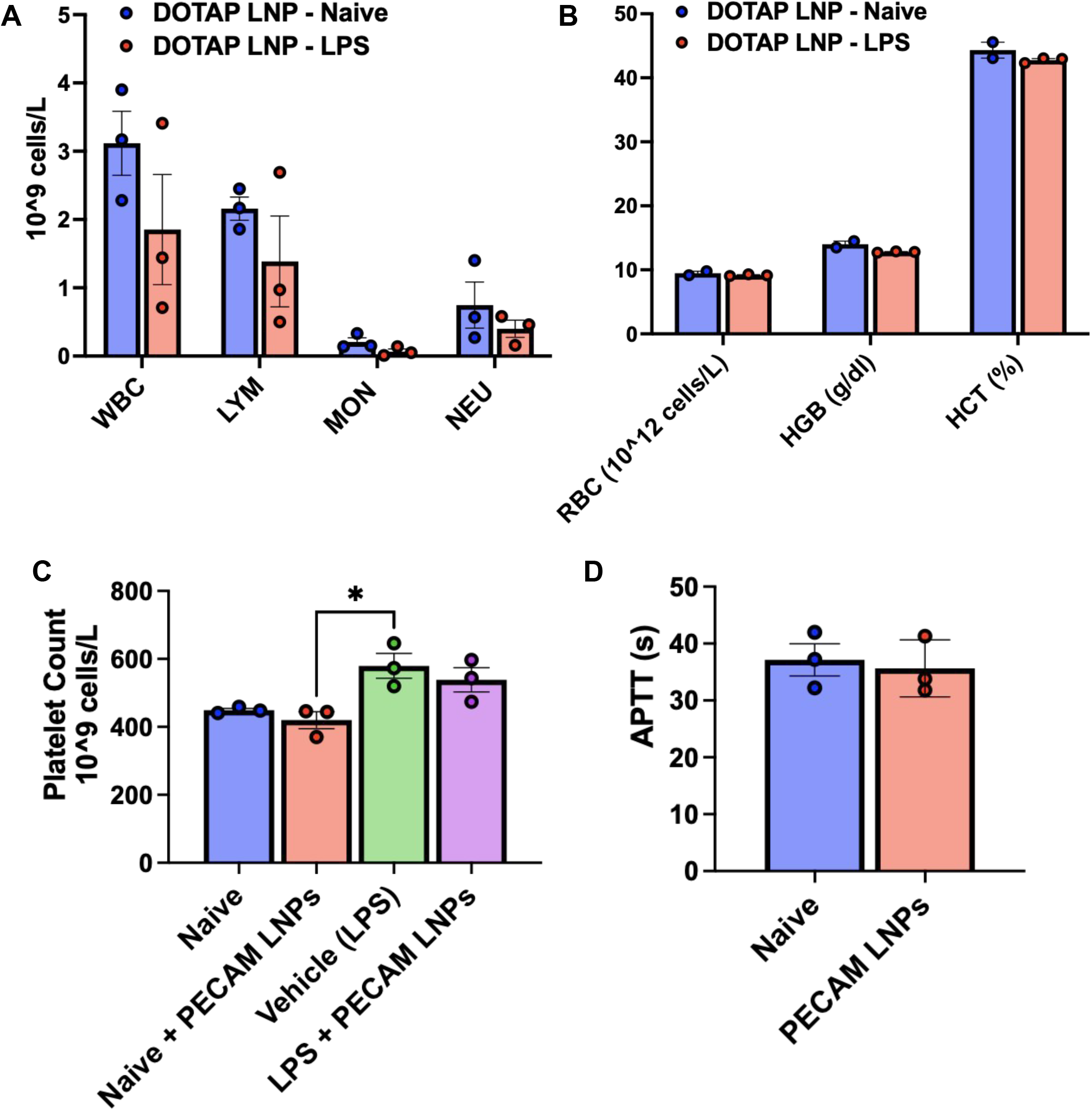
(A) Complete blood counts (CBCs) from naïve or nebulized-LPS mice injected with +DOTAP LNPs showing counts of white blood cells (WBC), lymphocytes (LYM), monocytes (MON), and neutrophils (NEU) (B) CBC showing counts of red blood cells (RBC), haemoglobin (HGB) and hematocrit (HCT) in naïve of nebulized-LPS mice injected with +DOTAP LNPs. (C) -DOTAP LNPs conjugated to antibodies against Platelet Endothelial Cell Adhesion Molecule (PECAM; a lung targeting moiety) do not affect platelet count in naive and nebulized-LPS mice 30 minutes post-LNP injection (D) PECAM LNPs maintain APTT at naïve levels *in vitro*.

**Supplementary Fig 8:**
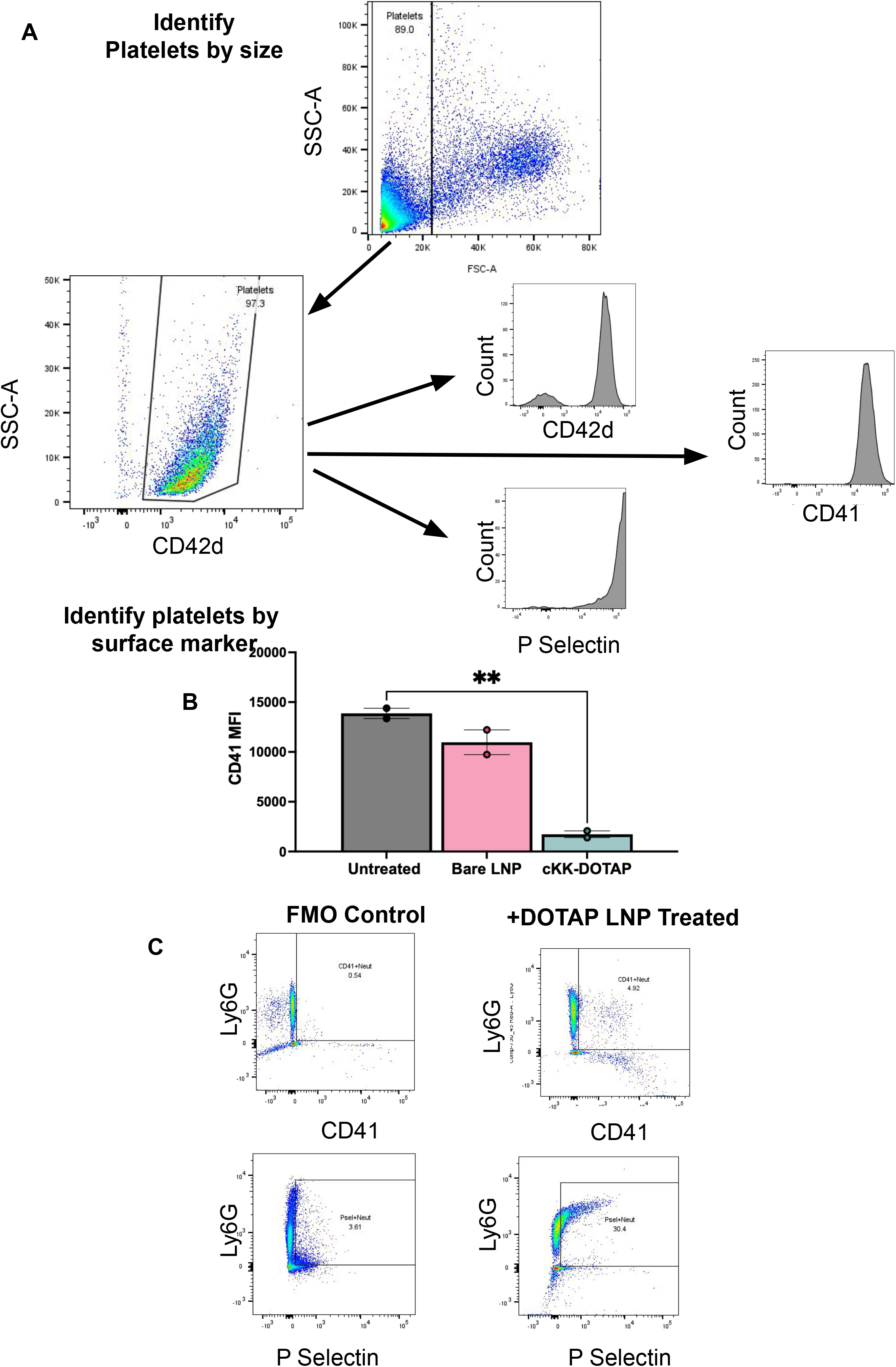
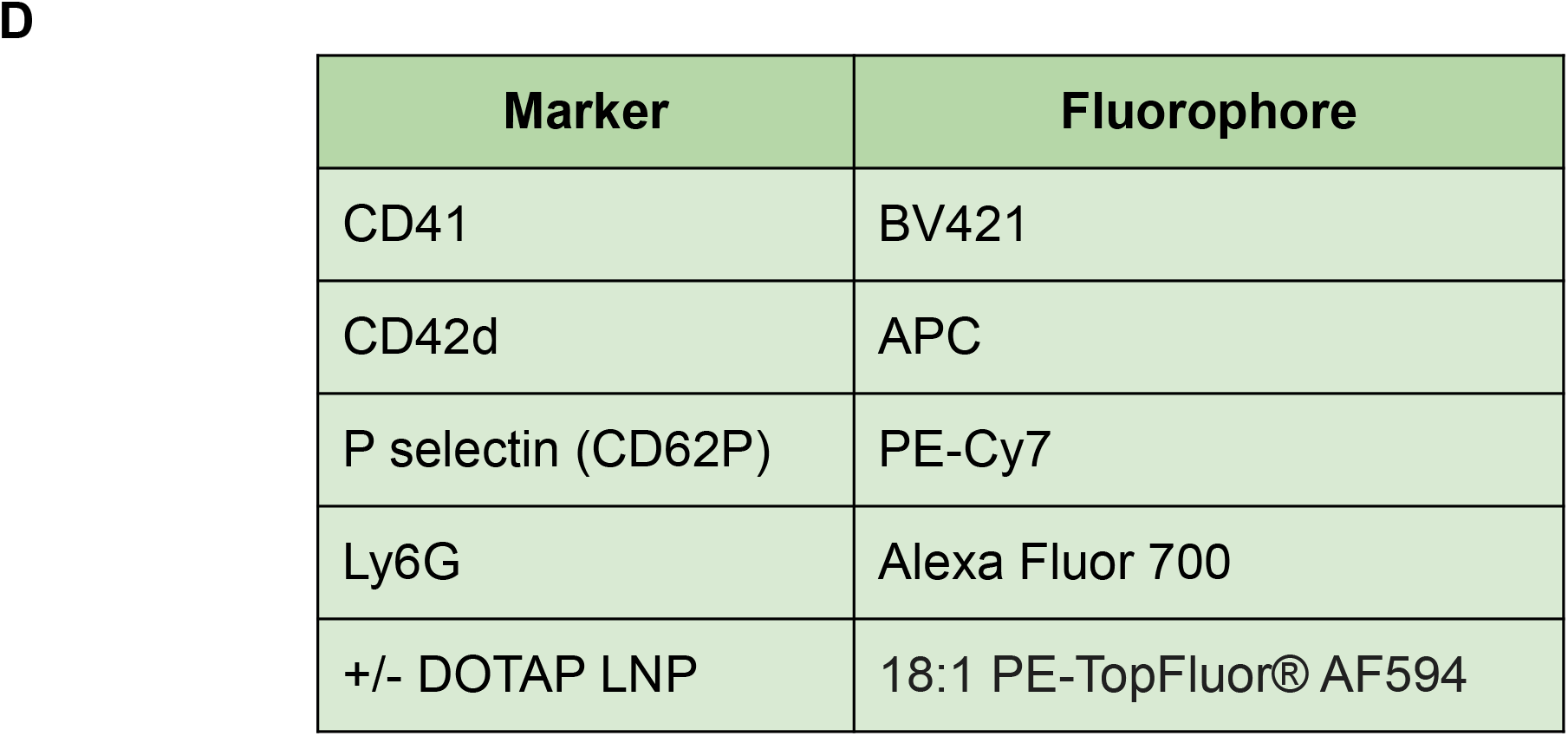
Flow cytometry gating for platelet flow cytometry and platelet-neutrophil aggregates. (A) Flow cytometry gating for platelet flow cytometry using a dual confirmation method, gating by both size of platelets as positivity for CD42d, a unique surface marker for platelets. (B)Flow cytometry MFI for CD41 expression on the surface of platelets shows a decrease with +DOTAP LNP treatment. (C) Flow cytometry plots showing that about 5% of neutrophils also stain positive for CD41 compared to fluorescence minus one (FMO) control. Similarly, about 30% of neutrophils also stain positive for p-selectin compared to FMO controls, showing the effects of thrombo-inflammation in the blood. (D) Table identifying the flow cytometry markers and the respective fluorophores used for flow cytometry experiments

**Supplementary Fig. 9:**
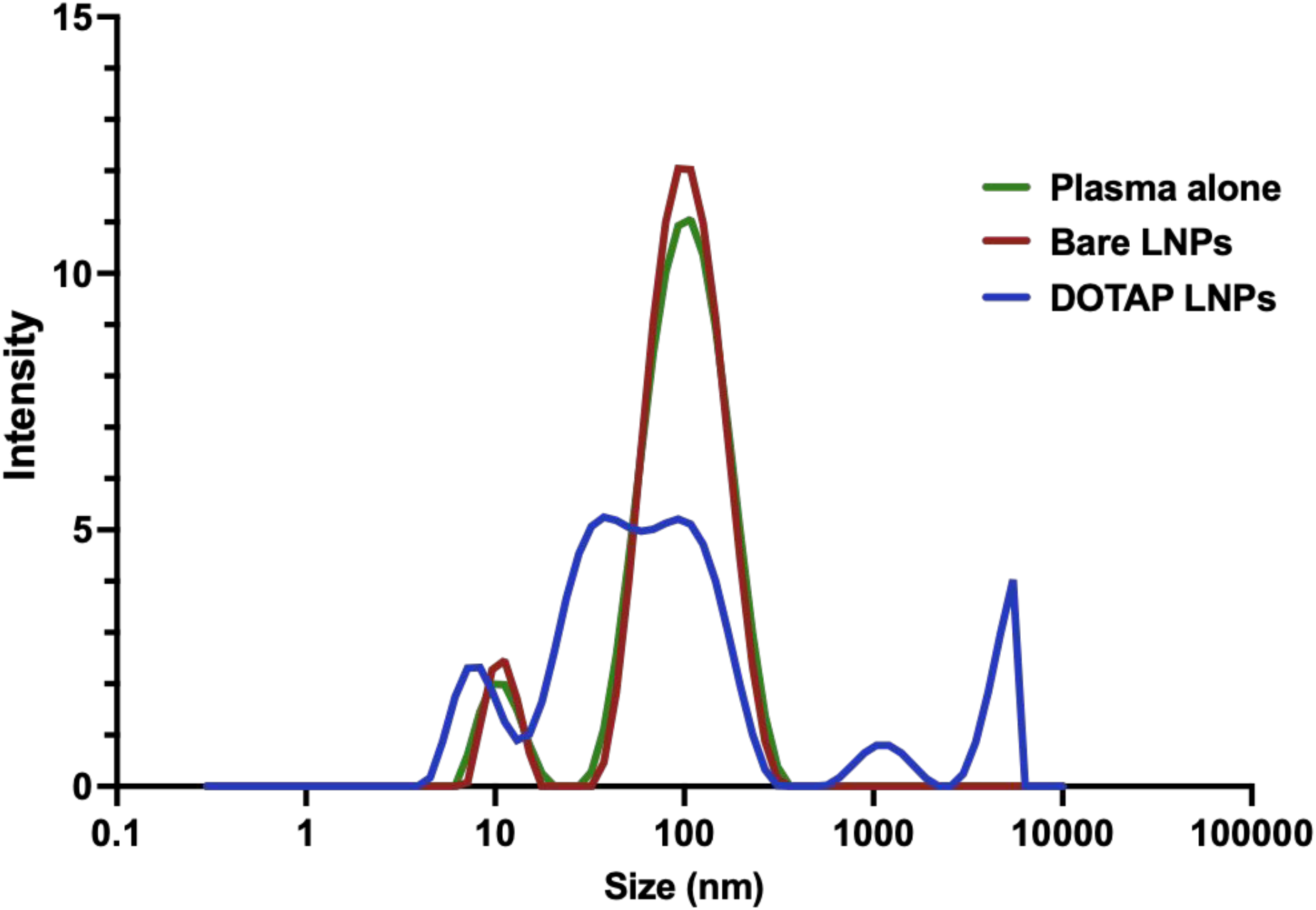
DLS size distributions of plasma alone or plasma treated with + or - DOTAP LNPs show aggregation with +DOTAP LNPs. Naïve mouse plasma was collected and incubated with LNPs at a dose of at 37℃ for 30 minutes. After a 100-fold dilution with PBS, the samples were read with a Zetasizer.

**Supplementary Fig. 10:**
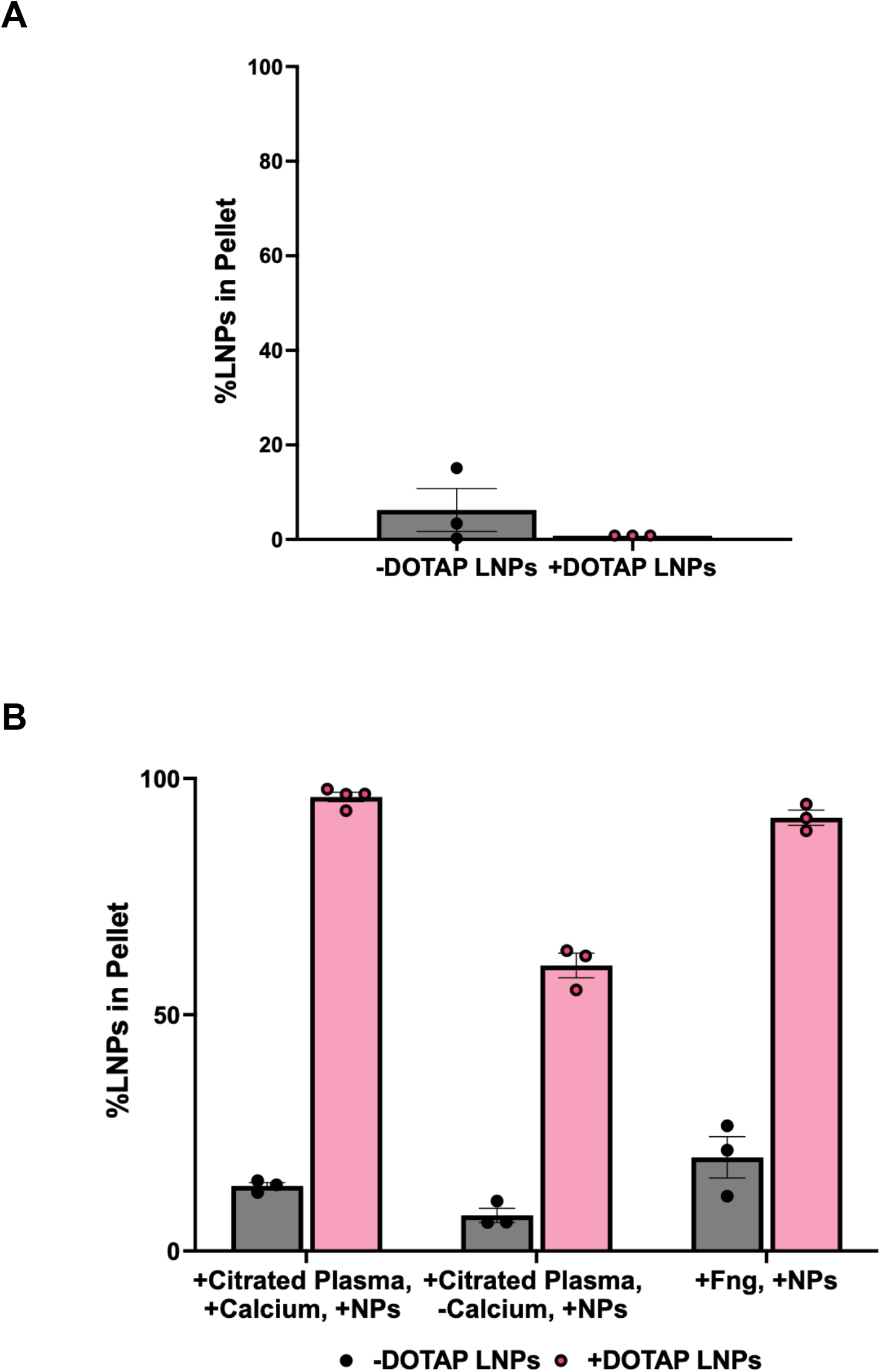
(A) - or + DOTAP LNPs do not precipitate significantly in PBS. (B) When incubated with citrated plasma, recalcified citrated plasma or fibrinogen alone, +DOTAP LNPs show significant aggregation and precipitation while -DOTAP LNPs do not. In these experiments, radioactively labeled LNPs were incubated in PBS, plasma or fibrinogen for 30 minutes at 37℃. After centrifugation for 10 minutes at 100 x g, the radioactivity in the pellet and the supernatant were quantified. The fraction of LNPs in the pellet can be interpreted as the fraction of LNPs that are aggregated.

**Supplementary Fig. 11:**
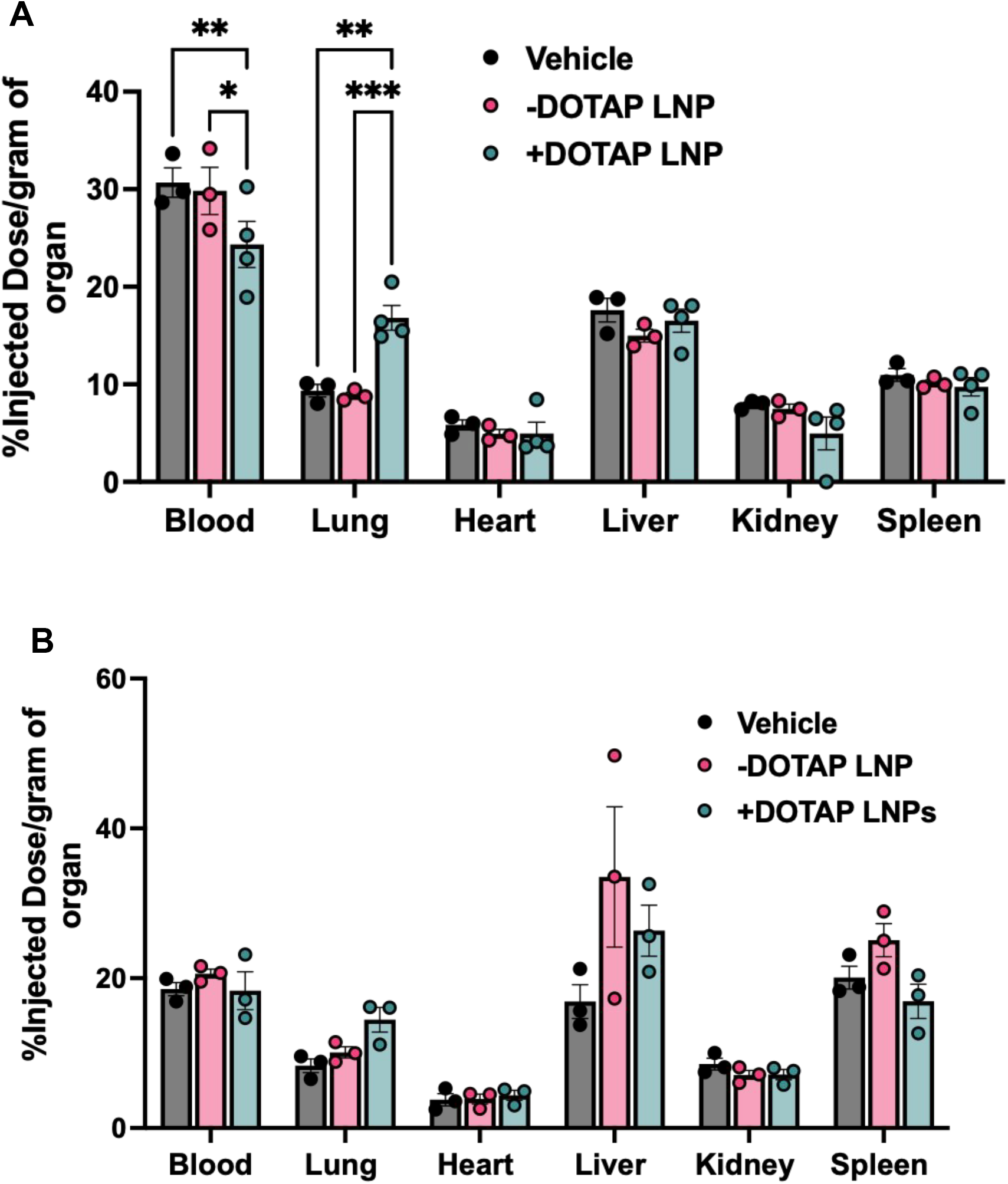
Biodistributions of fibrinogen in (A) naïve and (B) nebulized-LPS injured mice injected with fibrinogen alone or fibrinogen and + or - DOTAP LNPs.

**Supplementary Fig. 12:**
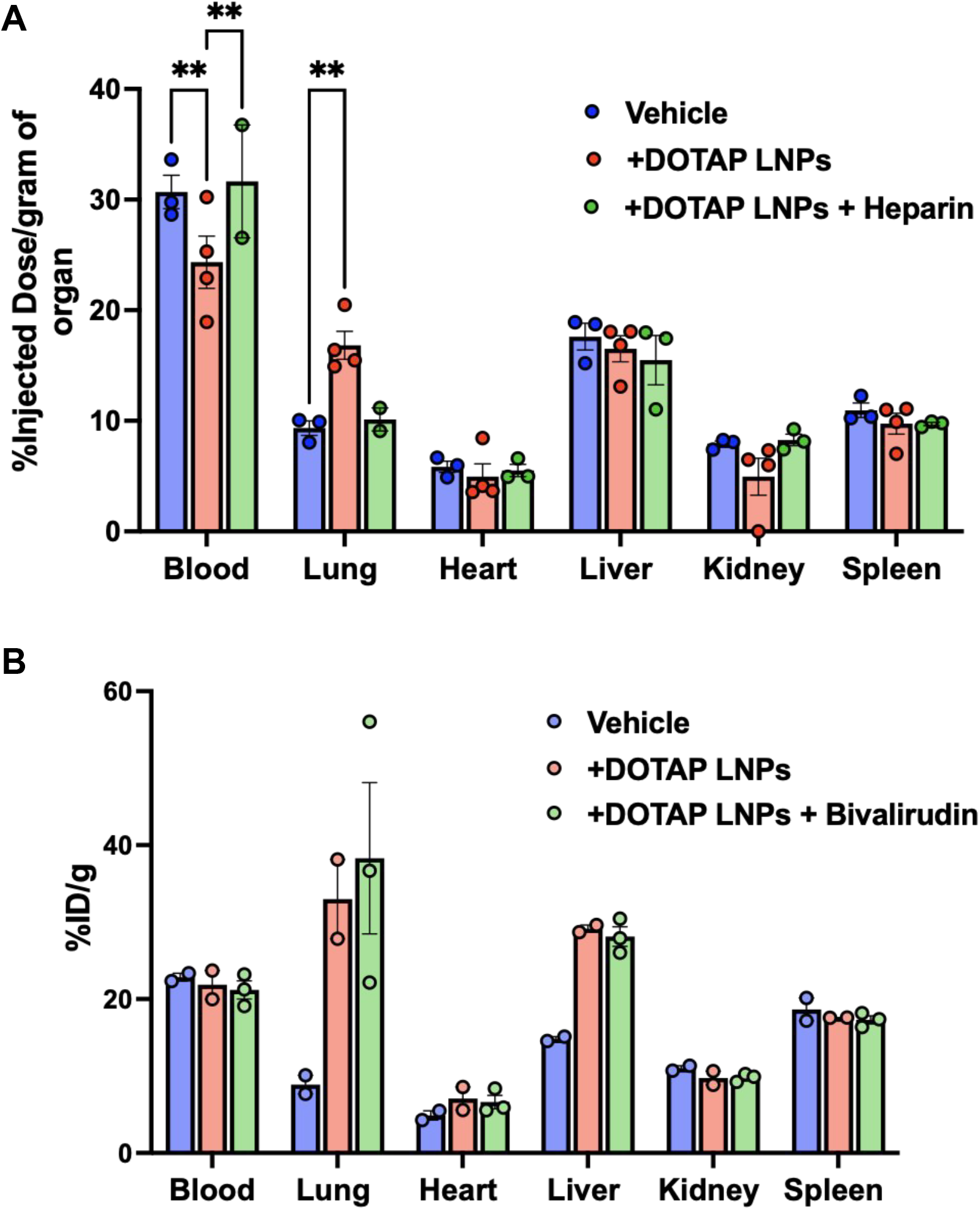
Biodistributions of fibrinogen after pre-treatment with the anticoagulants (A) heparin or (B) bivalirudin followed by +DOTAP LNP treatment. Heparin pre-treatment completely reverses fibrinogen lung uptake to vehicle levels while bivalirudin does not alter fibrinogen biodistribution compared to injection with +DOTAP LNPs alone.

**Supplementary Figure 13:**
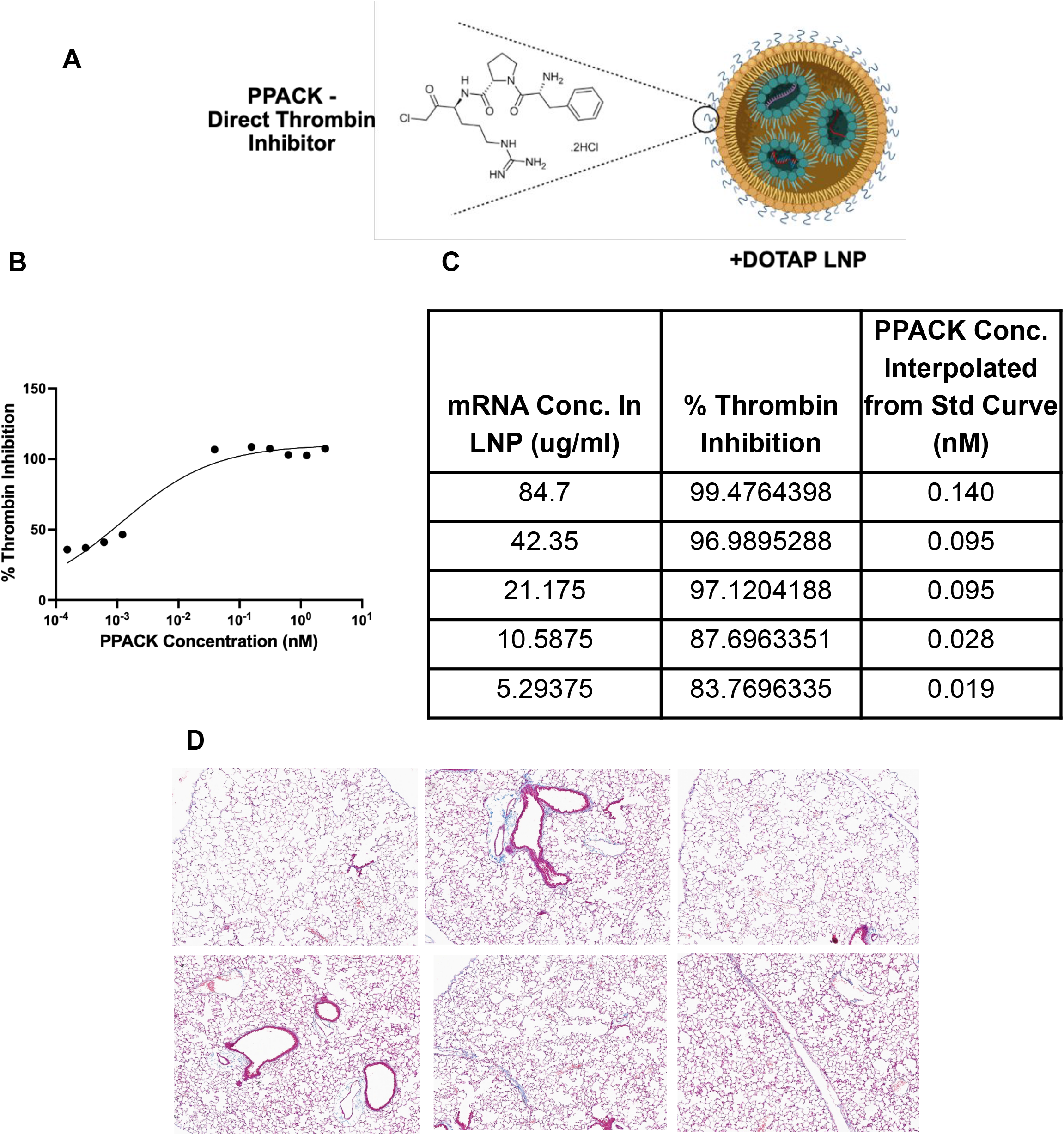
(A) The direct thrombin inhibitor PPACK was surface conjugated onto +DOTAP LNPs using azide-DBCO click chemistry. (B) A standard curve was generated to measure the inhibition of thrombin with PPACK at different concentrations using a Chromozym assay. PPACK was incubated with thrombin followed by the addition of the chromophoric thrombin substrate Chromozym. The change in absorbance at 405nm has a negative correlation with thrombin inhibition. (C) At undiluted concentrations, PPACK-conjugated +DOTAP LNPs have ∼100% inhibition of thrombin. (D) Lung histological samples from mice injected with +DOTAP LNPs show a significantly reduced appearance of clots. Unfortunately, these PPACK-LNPs were found to cause an increase in mouse mortality via an unknown mechanism, with our hypothesis being hemorrhage. Therefore, further engineering is needed to make anticoagulant-conjugated LNPs safe and effective.

## References

1. Chaudhary, N., Weissman, D. & Whitehead, K. A. mRNA vaccines for infectious diseases: principles, delivery and clinical translation. Nat. Rev. Drug Discov. 20, 817–838 (2021).

2. Parhiz, H. et al. PECAM-1 directed re-targeting of exogenous mRNA providing two orders of magnitude enhancement of vascular delivery and expression in lungs independent of apolipoprotein E-mediated uptake. J. Control. Release 291, 106–115 (2018).

3. Kim, M. et al. Engineered ionizable lipid nanoparticles for targeted delivery of RNA therapeutics into different types of cells in the liver. Sci Adv 7, (2021).

4. Akinc, A. et al. Targeted delivery of RNAi therapeutics with endogenous and exogenous ligand-based mechanisms. Mol. Ther. 18, 1357–1364 (2010).

5. Cheng, Q. et al. Selective organ targeting (SORT) nanoparticles for tissue-specific mRNA delivery and CRISPR--Cas gene editing. Nat. Nanotechnol. 15, 313–320 (2020).

6. Kranz, L. M. et al. Systemic RNA delivery to dendritic cells exploits antiviral defence for cancer immunotherapy. Nature 534, 396–401 (2016).

7. Zhang, D. et al. Targeted Delivery of mRNA with One-Component Ionizable Amphiphilic Janus Dendrimers. J. Am. Chem. Soc. 143, 17975–17982 (2021).

8. Dilliard, S. A., Cheng, Q. & Siegwart, D. J. On the mechanism of tissue-specific mRNA delivery by selective organ targeting nanoparticles. Proceedings of the National Academy of Sciences 118, e2109256118 (2021).

9. LoPresti, S. T., Arral, M. L., Chaudhary, N. & Whitehead, K. A. The replacement of helper lipids with charged alternatives in lipid nanoparticles facilitates targeted mRNA delivery to the spleen and lungs. J. Control. Release 345, 819–831 (2022).

10. Blake, T. R. et al. Lysine-Derived Charge-Altering Releasable Transporters: Targeted Delivery of mRNA and siRNA to the Lungs. Bioconjug. Chem. (2023) doi:10.1021/acs.bioconjchem.3c00019.

11. Kaczmarek, J. C. et al. Optimization of a Degradable Polymer–Lipid Nanoparticle for Potent Systemic Delivery of mRNA to the Lung Endothelium and Immune Cells. Nano Lett. 18, 6449–6454 (2018).

12. Yan, Y., Xiong, H., Zhang, X., Cheng, Q. & Siegwart, D. J. Systemic mRNA Delivery to the Lungs by Functional Polyester-based Carriers. Biomacromolecules 18, 4307–4315 (2017).

13. Myerson, J. W. et al. Supramolecular arrangement of protein in nanoparticle structures predicts nanoparticle tropism for neutrophils in acute lung inflammation. Nat. Nanotechnol. 17, 86–97 (2022).

14. Wang, Z. et al. Combating complement’s deleterious effects on nanomedicine by conjugating complement regulatory proteins to nanoparticles. Adv. Mater. 34, e2107070 (2022).

15. Fam, S. Y. et al. Stealth Coating of Nanoparticles in Drug-Delivery Systems. Nanomaterials (Basel) 10, (2020).

16. Wu, X. et al. Inhibition of intrinsic coagulation improves safety and tumor-targeted drug delivery of cationic solid lipid nanoparticles. Biomaterials 156, 77–87 (2018).

17. Ilinskaya, A. N. & Dobrovolskaia, M. A. Nanoparticles and the blood coagulation system. Part II: safety concerns. Nanomedicine 8, 969–981 (2013).

18. Jones, C. F. et al. Cationic PAMAM dendrimers disrupt key platelet functions. Mol. Pharm. 9, 1599–1611 (2012).

19. Oslakovic, C., Cedervall, T., Linse, S. & Dahlbäck, B. Polystyrene nanoparticles affecting blood coagulation. Nanomedicine 8, 981–986 (2012).

20. de la Harpe, K. M. et al. The Hemocompatibility of Nanoparticles: A Review of Cell– Nanoparticle Interactions and Hemostasis. Cells 8, 1209 (2019).

21. Fröhlich, E. Action of Nanoparticles on Platelet Activation and Plasmatic Coagulation. Curr. Med. Chem. 23, 408–430 (2016).

22. Maas, C. & Renné, T. Coagulation factor XII in thrombosis and inflammation. Blood 131, 1903–1909 (2018).

23. Esmon, C. T. Molecular circuits in thrombosis and inflammation. Thromb. Haemost. 109, 416–420 (2013).

24. Sang, Y., Roest, M., de Laat, B., de Groot, P. G. & Huskens, D. Interplay between platelets and coagulation. Blood Rev. 46, 100733 (2021).

25. Spurgeon, B. E. J., Linden, M. D., Michelson, A. D. & Frelinger, A. L., 3rd. Immunophenotypic Analysis of Platelets by Flow Cytometry. Curr Protoc 1, e178 (2021).

26. Blann, A. D. & Lip, G. Y. Hypothesis: is soluble P-selectin a new marker of platelet activation? Atherosclerosis 128, 135–138 (1997).

27. Bashiri, G. et al. Nanoparticle protein corona: from structure and function to therapeutic targeting. Lab Chip 23, 1432–1466 (2023).

28. Wang, S. et al. The role of protein corona on nanodrugs for organ-targeting and its prospects of application. J. Control. Release 360, 15–43 (2023).

29. Protopopova, A. D. et al. Visualization of fibrinogen αC regions and their arrangement during fibrin network formation by high-resolution AFM. J. Thromb. Haemost. 13, 570–579 (2015).

30. Jung, S.-Y. et al. The Vroman effect: a molecular level description of fibrinogen displacement. J. Am. Chem. Soc. 125, 12782–12786 (2003).

31. Derakhshankhah, H. et al. Molecular interaction of fibrinogen with zeolite nanoparticles. Sci. Rep. 9, 1558 (2019).

32. Deng, Z. J., Liang, M., Toth, I., Monteiro, M. J. & Minchin, R. F. Molecular interaction of poly(acrylic acid) gold nanoparticles with human fibrinogen. ACS Nano 6, 8962–8969 (2012).

33. Deng, Z. J., Liang, M., Monteiro, M., Toth, I. & Minchin, R. F. Nanoparticle-induced unfolding of fibrinogen promotes Mac-1 receptor activation and inflammation. Nat. Nanotechnol. 6, 39–44 (2011).

34. Savage, B., Saldívar, E. & Ruggeri, Z. M. Initiation of platelet adhesion by arrest onto fibrinogen or translocation on von Willebrand factor. Cell 84, 289–297 (1996).

35. Qiu, Y. et al. Platelet Mechanosensing: Adhesion and Spreading On Immobilized Fibrinogen Depends On Substrate Stiffness. Blood 120, 384–384 (2012).

36. Loncar, R. et al. Platelet adhesion onto immobilized fibrinogen under arterial and venous in-vitro flow conditions does not significantly differ between men and women. Thromb. J. 5, 5 (2007).

37. Mangin, P. H. et al. Immobilized fibrinogen activates human platelets through glycoprotein VI. Haematologica 103, 898–907 (2018).

38. Weiss, R. J., Esko, J. D. & Tor, Y. Targeting heparin and heparan sulfate protein interactions. Org. Biomol. Chem. 15, 5656–5668 (2017).

39. Myerson, J., He, L., Lanza, G., Tollefsen, D. & Wickline, S. Thrombin-inhibiting perfluorocarbon nanoparticles provide a novel strategy for the treatment and magnetic resonance imaging of acute thrombosis. J. Thromb. Haemost. 9, 1292–1300 (2011).

40. Parhiz, H. et al. Added to pre-existing inflammation, mRNA-lipid nanoparticles induce inflammation exacerbation (IE). J. Control. Release 344, 50–61 (2022).

41. Marcos-Contreras, O. A. et al. Selective targeting of nanomedicine to inflamed cerebral vasculature to enhance the blood-brain barrier. Proc. Natl. Acad. Sci. U. S. A. 117, 3405– 3414 (2020).

